# ProxiNet transfers spatially learned cellular proximity to dissociated single-cell transcriptomes

**DOI:** 10.64898/2026.09.25.754372

**Authors:** Yuxiang Zhan, Binwei Yan, Ao Zhang, Manolis Kellis, Na Sun

## Abstract

Spatial transcriptomics reveals cellular organization within intact tissues, whereas dissociated single-cell RNA sequencing provides broad transcriptomic coverage but loses information about cellular proximity and neighborhood structure. Here, we developed **ProxiNet**, a spatially supervised framework that learns transcriptomic signatures of pairwise cellular proximity from spatial reference datasets and transfers these relationships to dissociated single-cell transcriptomes. ProxiNet predicted cellular proximity across brain regions and spatial technologies, including zero-shot cross-technology transfer, and gradient-based attribution identified genes and broader transcriptional programs associated with proximity predictions. In spatial datasets with known coordinates, ProxiNet-derived cellular neighborhoods recovered reproducible tissue organization and anatomical structure, providing independent spatial validation of the inferred proximity relationships. Applying the spatially calibrated model to dissociated scRNA-seq revealed heterogeneous cellular neighborhoods with distinct cell-type compositions and candidate communication programs. In an Alzheimer’s disease dataset, ProxiNet further identified age-associated remodeling of inferred neighborhoods, including an AD-associated neighborhood at 8 months characterized by distinct astrocyte and neuronal transcriptional states and candidate intercellular communication programs. Together, these results establish pairwise cellular proximity as an interpretable and transferable representation for extending spatially learned tissue organization to dissociated single-cell transcriptomes.

## Introduction

Single-cell RNA sequencing has transformed our ability to define cellular diversity and molecular states across tissues, development, and disease^1^. However, tissue dissociation removes information about where cells were located, and which cells occupied shared local environments. This loss is biologically important because cell state and function depend not only on intrinsic molecular identity but also on interactions with neighboring cells and local tissue architecture, which can define functionally distinct tissue units and reveal biological organization that is not apparent from cell-type abundance alone^2,3^.

Spatial transcriptomic technologies address this limitation by measuring molecular profiles in intact tissue, but current platforms involve trade-offs among gene coverage, spatial resolution, and throughput. Imaging-based approaches such as MERFISH and STARmap provide cellular spatial resolution but typically measure a targeted gene set, whereas sequencing-based methods such as Slide-seq and 10x Visium provide broader transcriptomic coverage without subcellular resolution. These complementary strengths have motivated computational approaches that integrate spatial and dissociated transcriptomes^3^. Methods such as Tangram and CytoSPACE map dissociated cells onto spatial references, while novoSpaRc and CeLEry infer spatial positions or origins from gene expression^4–7^. These approaches are particularly useful when reconstructing spatial coordinates or assigning cells to measured tissue locations is the primary goal.

More recent methods have shifted from coordinate reconstruction toward learning local spatial context. CellContrast uses spatially supervised contrastive learning to generate representations in which spatially proximal cells have similar embeddings and uses these representations to reconstruct spatial relationships in dissociated cells^8^. ENVI jointly models spatial and dissociated transcriptomes and transfers a covariance-based representation of cellular niches to scRNA-seq^9^. Nicheformer further demonstrates that large-scale joint training on spatial and dissociated cells can predict spatial labels and neighborhood composition from dissociated transcriptomes^10^. Together, these studies show that transcriptional state contains information about cellular spatial context. However, these frameworks primarily represent spatial context through reconstructed positions, latent similarities, niche representations, or predefined spatial properties, and evaluate the performance on a global scale. An explicit prediction of pairwise cellular proximity provides a complementary relational representation at the single-cell scale that can be directly assembled into a cell graph and interpreted at the level of individual cell pairs.

Here, we developed ProxiNet, a spatially supervised graph-learning framework that treats pairwise cellular proximity as the prediction target. ProxiNet uses spatial coordinates to define proximal and non-proximal cell pairs during training, while representing transcriptomic information through a heterogeneous cell-gene graph. Once trained, the model predicts pairwise proximity probabilities from dissociated transcriptomes without requiring spatial coordinates at inference. We found that proximity signatures generalized across brain regions and across MERFISH, STARmap, and Slide-seq, including zero-shot cross-technology transfer and further improvement after fine-tuning. We further used gradient-based attribution to identify proximity-associated genes and connect these genes to broader transcriptomic programs. Predicted proximity graphs recovered tissue organization in spatial datasets and revealed heterogeneous cellular neighborhoods in dissociated scRNA-seq. Finally, applying ProxiNet to an Alzheimer’s disease dataset identified age-associated remodeling of inferred cellular neighborhoods and neighborhood-associated transcriptional programs. Together, these analyses establish pairwise cellular proximity as an interpretable and transferable representation linking spatial atlases to dissociated single-cell transcriptomes.

## Results

### ProxiNet learns transferable transcriptomic signatures of cellular proximity

Dissociated single-cell RNA sequencing provides detailed molecular profiles of individual cells but removes information about which cells occupied nearby positions in the tissue. We reasoned that pairwise cellular proximity could provide a transferable representation of tissue organization that can be learned from spatial transcriptomic data and later inferred from dissociated transcriptomes. We therefore developed **ProxiNet**, a spatially supervised framework that learns transcriptomic features associated with cellular proximity. We trained and evaluated ProxiNet using mouse whole-brain spatial transcriptomic datasets generated by three technologies, MERFISH^11^, STARmap^12^, and Slide-seq^13^, with spatial sections assigned to training, validation, and test sets (**Supplementary Table 1-3, Methods**). During spatial calibration, we used measured spatial coordinates to label pairs of cells as proximal or non-proximal according to a predefined distance threshold and trained ProxiNet to predict these labels from gene expression. At inference, ProxiNet requires only transcriptomic profiles and generates a probability of proximity for each cell pair. We then used these probabilities to construct a weighted cellular-proximity graph that can be further partitioned into inferred cellular neighborhoods (**Fig. 1a, Methods**). This formulation allows ProxiNet to transfer information about relative cellular organization without requiring reconstruction of absolute spatial coordinates.

**Figure 1.**
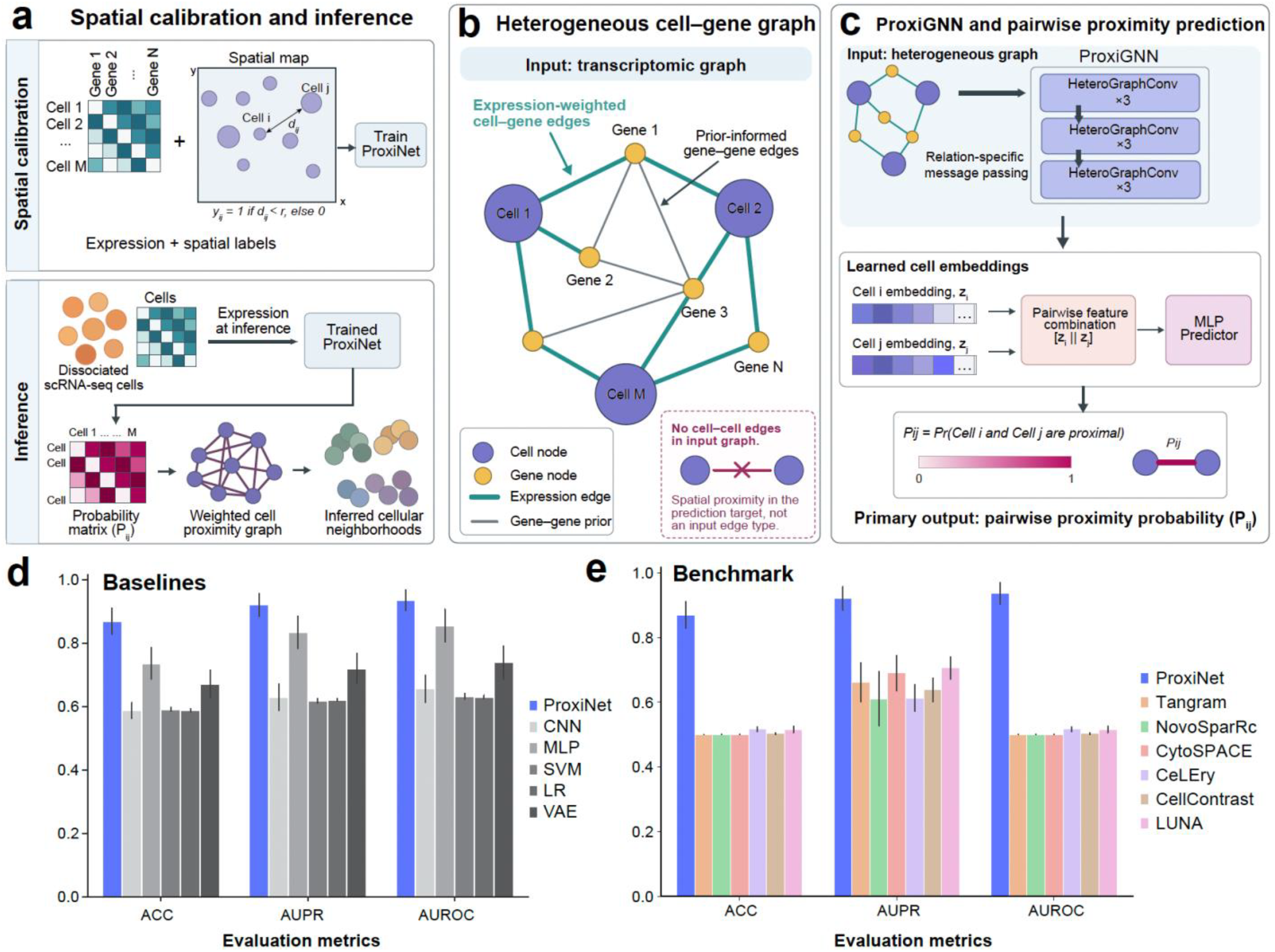
ProxiNet learns transferable transcriptomic signatures of cellular proximity. **a**, Overview of spatial calibration and inference with ProxiNet. During spatial calibration, gene-expression profiles are paired with measured spatial coordinates, and pairs of cells are labeled as proximal when their spatial distance d_*ij*_ is below a predefined distance threshold *r*. ProxiNet is trained to predict these pairwise proximity labels from transcriptomic information. At inference, the trained model is applied to dissociated scRNA-seq cells using gene expression alone to generate a pairwise proximity-probability matrix *P*_*ij*_. Predicted probabilities define a weighted cellular-proximity graph that can subsequently be partitioned into inferred cellular neighborhoods. **b**, Heterogeneous transcriptomic graph used as input to ProxiNet. Cell and gene nodes are connected by expression-weighted cell–gene edges, while gene nodes are connected by prior-informed gene–gene edges. No cell–cell edges are provided as model inputs; cellular proximity is used only as the spatially derived prediction target during training. **c**, ProxiNet architecture for pairwise proximity prediction. Relation-specific message passing with ProxiGNN generates cell embeddings ***z***_*i*_ and ***z***_*j*_, which are combined and passed to an MLP predictor to estimate *P*_*ij*_ = *Pr(cells i and cell j are proximal)*. **d**, Comparison of ProxiNet with CNN, MLP, support vector machine (SVM), logistic regression (LR), and variational autoencoder (VAE) baselines using accuracy (ACC), area under the precision–recall curve (AUPR), and area under the receiver operating characteristic curve (AUROC). **e**, Comparison of ProxiNet with Tangram, NovoSpaRc, CytoSPACE, CeLEry, CellContrast, and LUNA using the same proximity-prediction evaluation metrics. Bars show the value of evaluation metrics and error bars indicate standard deviation across sections in test data.

To learn proximity-associated representations from transcriptomic data, we represented each dataset as a heterogeneous cell-gene graph. We connected cell nodes to gene nodes using expression-weighted edges and connected gene nodes using prior-informed gene-gene relationships (**Fig. 1b, Supplementary Table 4, Methods**). Importantly, we did not provide cell-cell edges to the model. Instead, we used spatial proximity only as the prediction task during training. ProxiGNN performs relation-specific message passing across cell-to-gene, gene-to-cell, and gene-to-gene relations and generates an embedding for each cell. The pairwise predictor combines the embeddings of two cells and estimates the probability that they are spatially proximal (**Fig. 1c, Methods**). The detailed architecture uses stacked heterogeneous graph convolution layers, followed by an MLP for pairwise proximity prediction (**Supplementary Fig. 1a-c**). Thus, ProxiNet learns cellular proximity from transcriptomic relationships rather than encoding known cellular adjacency directly into the input graph.

We first evaluated whether this graph-based formulation improved pairwise proximity prediction relative to conventional machine-learning approaches. Across test sections, ProxiNet achieved higher accuracy, area under the precision-recall curve, and area under the receiver operating characteristic curve than CNN^14^, MLP^15^, SVM^16^, logistic regression, and VAE^17^ baselines (**Fig. 1d**). We also compared ProxiNet with existing computational methods developed for spatial mapping, spatial reconstruction, or spatial representation learning, including Tangram^5^, NovoSpaRc^4^, CytoSPACE^7^, CeLEry^6^, CellContrast^8^, and LUNA^18^. We converted the outputs of these methods into a common pairwise-proximity evaluation framework and found that ProxiNet achieved higher performance across all three proximity-prediction metrics (**Fig. 1e**). These results indicate that directly optimizing pairwise cellular proximity provides a more effective representation for this task than either conventional transcriptomic models or methods designed primarily to recover spatial coordinates.

We next asked whether ProxiNet learned dataset-specific spatial patterns or more transferable transcriptomic signatures of proximity. Using the mouse whole-brain spatial datasets described above, we designed three train, validation, and test configurations in which anatomical regions were held out for evaluation (**Supplementary Fig. 1d-f, Methods**). We found that ProxiNet maintained similar AUPR and AUROC across held-out anatomical regions. We then evaluated performance separately across MERFISH, STARmap, and Slide-seq datasets and found consistently high proximity-prediction performance across all three technologies (**Supplementary Fig. 1g**). In cross-technology experiments, we first evaluated zero-shot transfer by applying models trained on one spatial technology directly to another without additional training. As expected, performance was highest when training and testing used the same platform, but ProxiNet retained substantial predictive ability across technologies. For example, a model trained on MERFISH achieved an AUPR of 0.72 on STARmap and 0.63 on Slide-seq, compared with 0.92 on MERFISH, while the corresponding AUROC values were 0.75, 0.60, and 0.94, respectively (**Supplementary Fig. 1h**). We observed similar zero-shot transfer for models trained on STARmap and Slide-seq. We next fine-tuned each pretrained model using data from the target technology and found that cross-technology performance increased substantially, with AUPR values reaching 0.89 to 0.93 and AUROC values reaching 0.91 to 0.94 across most train-test combinations (**Supplementary Fig. 1i**). These results suggest that ProxiNet learns transcriptomic signatures of cellular proximity that are transferable across measurement technologies and can be further adapted efficiently to a new spatial platform through fine-tuning.

Finally, we asked whether the learned representations retained additional information about spatial organization beyond the supervised pairwise proximity task. As an auxiliary analysis, we used ProxiNet representations for reference-based coordinate mapping and direct coordinate inference and compared their performance with established spatial mapping and reconstruction methods (**Supplementary Fig. 2a, Supplementary Table 5, Methods**). We found that ProxiNet representations supported recovery of spatial organization across multiple independent sections, with competitive Jensen-Shannon divergence and Spearman correlation relative to Tangram, NovoSpaRc, CytoSPACE, CeLEry, CellContrast, and LUNA (**Supplementary Fig. 2b-e**). These results are consistent with the idea that supervision on pairwise proximity induces representations that retain broader information about tissue organization, even though coordinate reconstruction is not the primary objective of ProxiNet. Together, these analyses establish that ProxiNet learns transferable transcriptomic features of cellular proximity that generalize across datasets, brain regions, and spatial technologies.

### ProxiNet identifies transcriptomic programs associated with cellular proximity

We next asked which transcriptomic features contributed most strongly to ProxiNet proximity predictions. Because ProxiNet estimates proximity directly from gene-expression-derived cell representations, we used gradient-based attribution to trace each predicted proximity score back through the pairwise predictor and ProxiGNN to the expression-weighted cell-gene edges (**Fig. 2a, Methods**)^19^. This analysis produced a gene-level attribution score for each cell in a predicted cell pair. We refer to genes with high attribution scores as proximity-associated genes because they contribute strongly to the model prediction, although these scores do not imply that the genes causally determine cellular proximity. This framework allowed us to move beyond prediction accuracy and ask whether ProxiNet learned biologically structured transcriptomic features associated with cells that occupy nearby spatial environments.

**Figure 2.**
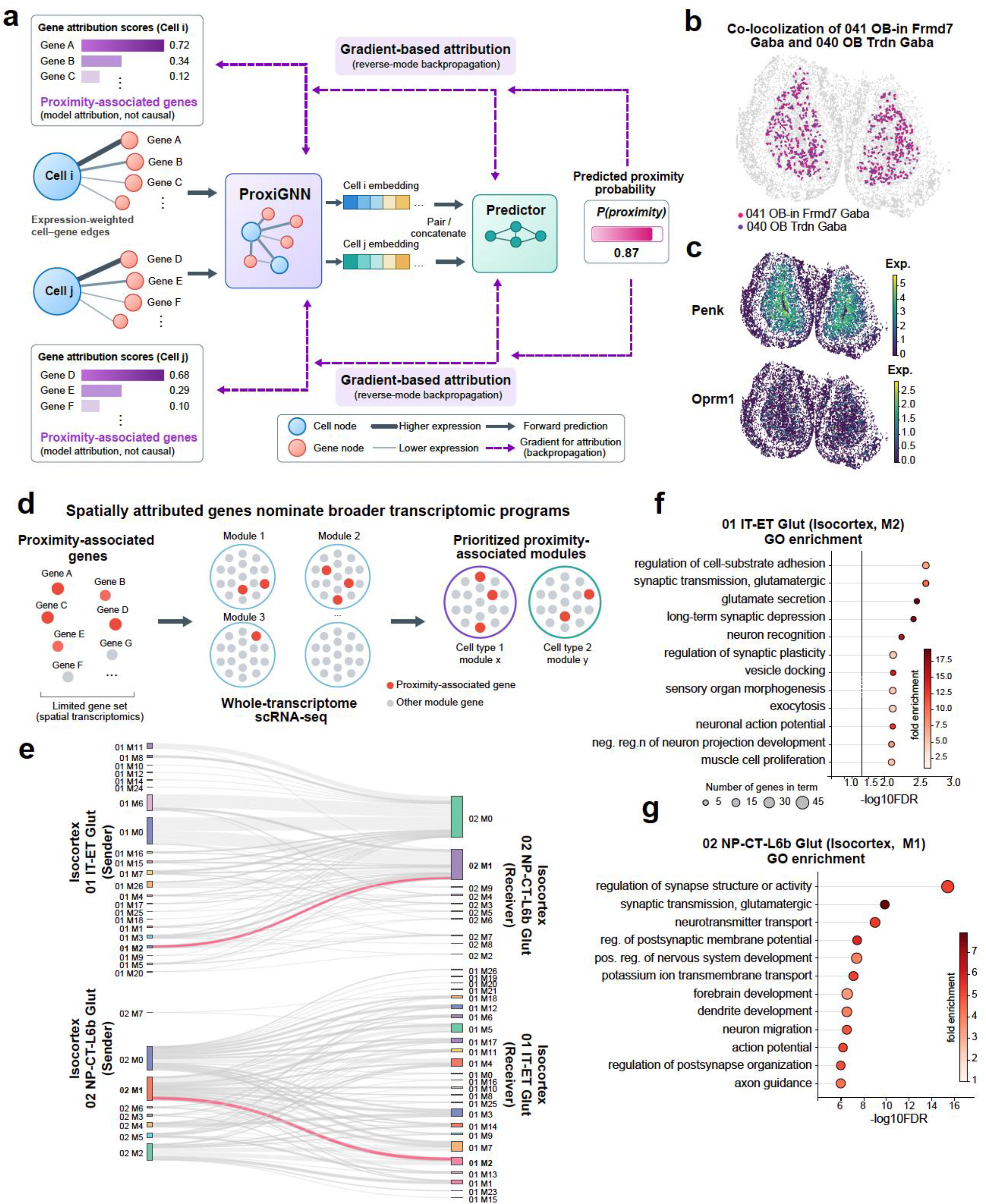
ProxiNet identifies transcriptomic programs associated with cellular proximity. **a**, Gradient-based attribution of ProxiNet proximity predictions to genes. Transcriptomic information from two cells is propagated through ProxiGNN to generate cell embeddings, which are combined by the pairwise predictor to estimate cellular-proximity probability. Reverse-mode gradient attribution traces the prediction back through the model to expression-weighted cell–gene edges, yielding gene-level attribution scores separately for each cell. Genes with high attribution are referred to as proximity-associated genes; these scores represent model attribution and do not imply causal effects on cellular proximity. **b**, Spatial distribution of the 041 OB-in Frmd7 GABA and 040 OB Trdn GABA cell populations in the olfactory bulb, illustrating their spatial co-localization. **c**, Spatial expression of *Penk* and *Oprm1* in the same tissue, providing an example of a ligand–receptor pair prioritized from proximity-associated genes in spatially co-localized cell populations. **d**, Strategy for extending proximity-associated genes identified from the spatial-transcriptomics gene panel to broader transcriptomic programs. Spatially attributed genes are mapped to modules derived from whole-transcriptome scRNA-seq, enabling prioritization of proximity-associated modules in defined cell types. **e**, Relationships between transcriptomic modules linked by ligand and receptor genes associated with sender and receiver states in Isocortex 01 IT–ET Glut and 02 NP–CT–L6b Glut cells. Highlighted connections denote ligand-receptor pairs which are significantly enriched in MERFISH dataset. **f**, Gene Ontology enrichment of proximity-associated module M2 in Isocortex 01 IT–ET Glut cells. **g**, Gene Ontology enrichment of proximity-associated module M1 in Isocortex 02 NP–CT–L6b Glut cells. Dot position denotes −log_10_FDR, dot color denotes fold enrichment, and dot size denotes the number of genes in the term.

We first tested whether highly attributed genes were enriched for known ligand-receptor relationships^20,21^. Across sender and receiver analyses, we observed a strong concentration of low enrichment P values using Fisher’s exact test, suggesting that ligand-receptor genes were preferentially represented among genes with high proximity attribution scores (**Supplementary Fig. 3a, Methods**). We then examined individual cell-type pairs in greater detail. For 041 OB-in Frmd7 GABA and 040 OB Trdn GABA cells, ligand-receptor genes were significantly enriched among the top attributed genes for both cell populations (**Supplementary Fig. 3b**). We found a similar pattern for 061 STR D1 GABA and 062 STR D2 GABA cells (**Supplementary Fig. 3c**). These results suggest that the transcriptomic features prioritized by ProxiNet are enriched for genes with potential roles in intercellular communication, consistent with the expectation that spatially proximal cells can share molecular programs associated with their local cellular environment.

We next examined whether individual proximity-associated genes corresponded to spatially coherent cell populations and candidate ligand-receptor relationships. In the olfactory bulb, we found that 041 OB-in Frmd7 GABA and 040 OB Trdn GABA cells occupied overlapping spatial domains (**Fig. 2b**). Among the highly attributed genes for these populations, *Penk* and *Oprm1* formed a candidate ligand-receptor pair and showed spatial expression patterns that overlapped with the corresponding cell populations (**Fig. 2c**). We observed an independent example in the striatum, where 061 STR D1 GABA and 062 STR D2 GABA cells co-localized spatially and the proximity attribution analysis prioritized the *Penk-Oprd1* pair (**Supplementary Fig. 3d**). These examples are consistent with the global enrichment analysis and indicate that ProxiNet attribution can nominate molecular features associated with spatially co-localized cell populations.

Because targeted spatial transcriptomic assays such as MERFISH measure only a subset of the transcriptome, we next asked whether spatially identified proximity-associated genes could be used to prioritize broader transcriptional programs in whole-transcriptome scRNA-seq^22^. We first identified gene modules from scRNA-seq per cell type per brain region from the whole brain dataset and then used proximity-associated genes measured in the spatial dataset to nominate modules containing these genes (**Fig. 2d, Methods**). This strategy allowed us to extend spatially calibrated attribution signals from a limited spatial gene panel to broader transcriptomic programs. In Isocortex, we examined module relationships between 01 IT-ET Glut and 02 NP-CT-L6b Glut cells in both sender and receiver directions. We found multiple module pairs connected by ligand-receptor genes, and several of these connections were significantly enriched in the MERFISH data (**Fig. 2e**). Gene Ontology analysis showed that the highlighted 01 IT-ET Glut module M2 was enriched for processes related to glutamatergic synaptic transmission, vesicle docking, synaptic plasticity, exocytosis, and neuronal activity (**Fig. 2f**). The highlighted 02 NP-CT-L6b Glut module M1 was enriched for synapse organization, glutamatergic transmission, neurotransmitter transport, dendrite development, neuronal migration, and axon guidance (**Fig. 2g**). We therefore found that spatially attributed genes nominate broader modules with coherent neuronal functions, suggesting that proximity-associated signals captured from targeted spatial measurements can identify more extensive transcriptomic programs in dissociated scRNA-seq.

We observed the same pattern in an independent analysis in another brain region, striatum. Proximity-associated modules connected 09 CNU-LGE GABA and 30 Astro-Epen cells in both sender and receiver directions (**Supplementary Fig. 3e**). We found that the prioritized CNU-LGE GABA module was enriched for neuronal functions including action potential, learning or memory, regulation of synapse organization, postsynaptic membrane potential, axon guidance, and calcium ion transport. The corresponding Astro-Epen module was enriched for programs including calcium-mediated signaling, ERK1 and ERK2 signaling, canonical Wnt signaling, neuron migration, telencephalon development, and cell projection organization (**Supplementary Fig. 3f**). These findings are consistent with the Isocortex results and suggest that ProxiNet attribution can connect spatially learned proximity signals to broader, cell-type-specific transcriptional programs across distinct brain regions. Together, these analyses show that ProxiNet not only predicts cellular proximity but also identifies genes and transcriptomic modules associated with those predictions.

### Inferred cellular neighborhoods recover spatial tissue organization

We next asked whether the pairwise proximity relationships predicted by ProxiNet could be assembled into higher-order cellular structures that correspond to tissue organization. We represented cells as nodes in a weighted cellular-proximity graph, with edge weights defined by ProxiNet-predicted pairwise proximity probabilities, and applied graph clustering to identify groups of cells with similar proximity relationships (**Fig. 3a, Methods**). We refer to these groups as inferred cellular neighborhoods. This formulation differs from coordinate reconstruction because it does not assign an absolute spatial position to each cell. Instead, it organizes cells according to their predicted local relationships with other cells. We therefore used spatial transcriptomic datasets with measured coordinates to test whether the inferred neighborhoods corresponded to recognizable tissue organization.

**Figure 3.**
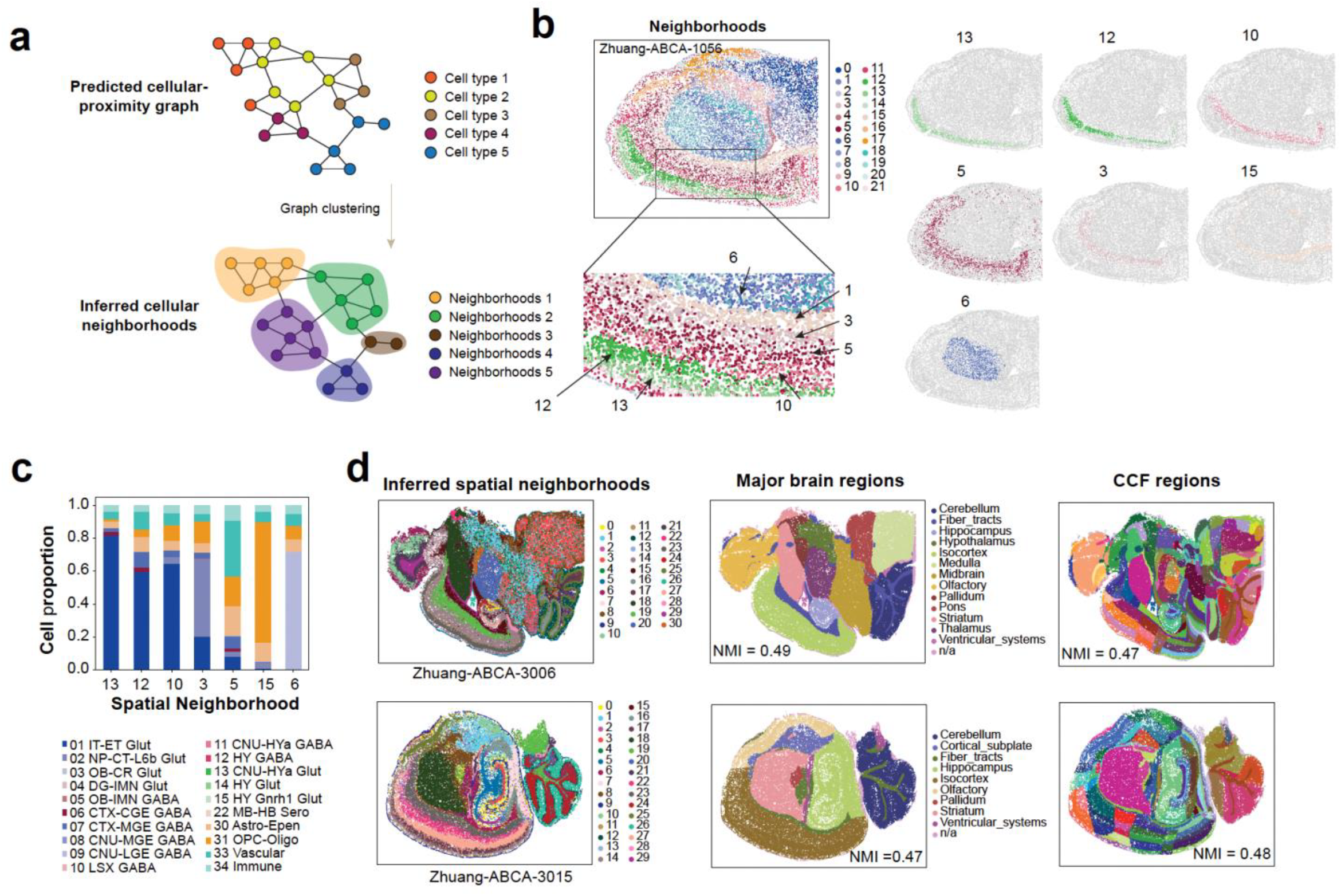
Inferred cellular neighborhoods recover spatial tissue organization. **a**, Construction of inferred cellular neighborhoods from ProxiNet predictions. Pairwise proximity probabilities define a weighted cellular - proximity graph, which is partitioned by graph clustering to identify groups of cells with similar inferred neighborhood organization. **b**, Spatial validation of inferred neighborhoods in the Zhuang-ABCA-1056 spatial transcriptomic dataset. Left, cells colored by inferred neighborhoods across the tissue section, with an enlarged region illustrating local organization of selected neighborhoods. Right, spatial distributions of representative neighborhoods 13, 12, 10, 5, 3, 15, and 6 shown individually. **c**, Cell-type composition of the selected inferred neighborhoods from panel B. Bars show the proportion of major neuronal and non-neuronal cell classes within each neighborhood. **d**, Comparison of inferred cellular neighborhoods with anatomical organization in two independent spatial datasets, Zhuang-ABCA-3006 and Zhuang-ABCA-3015. Cells are shown according to inferred neighborhood assignments, annotated major brain regions, and Common Coordinate Framework (CCF) regions. Concordance quantified by normalized mutual information (NMI) was 0.49 and 0.47 for inferred neighborhoods versus major brain regions and CCF regions, respectively, in Zhuang-ABCA-3006, and 0.47 and 0.48, respectively, in Zhuang-ABCA-3015.

In the Zhuang-ABCA-1056 dataset, we found that inferred cellular neighborhoods occupied spatially coherent regions within the tissue section (**Fig. 3b**). Within the cortical region, several neighborhoods formed adjacent and spatially ordered domains, and the individual distributions of neighborhoods 13, 12, 10, 5, 3, 15, and 6 showed a laminar organization consistent with the known layered architecture of the cerebral cortex^11^. We then examined the cellular composition of these neighborhoods and found that they contained distinct mixtures of neuronal and non-neuronal cell classes (**Fig. 3c**). These results indicate that clustering the predicted proximity graph recovers structured multicellular organization that reflects known tissue architecture rather than simply grouping cells according to a single dominant cell type.

We next asked whether this organization corresponded to established anatomical structure. In the independent Zhuang-ABCA-3006 dataset, inferred cellular neighborhoods showed concordance with both major brain-region annotations and Common Coordinate Framework regions, with normalized mutual information values of 0.49 and 0.47, respectively (**Fig. 3d**). We found a similar pattern in Zhuang-ABCA-3015, with NMI values of 0.47 for major brain regions and 0.48 for CCF regions. These moderate but reproducible levels of concordance indicate that the inferred neighborhoods capture substantial anatomical organization with detailed layer distinction while not simply reproducing the predefined anatomical labels. This is consistent with the idea that ProxiNet organizes cells according to learned local cellular relationships that reflect, but are not identical to, conventional brain-region boundaries.

We observed similar results across additional spatial datasets. In Zhuang-ABCA-1056 and Zhuang-ABCA-2031, inferred cellular neighborhoods showed NMI values of 0.41 and 0.40 with major brain-region annotations and 0.48 and 0.43 with CCF regions, respectively (**Supplementary Fig. 4a**). In the larger Zhuang-ABCA-3001 and Zhuang-ABCA-3008 sections, we found NMI values of 0.48 with major brain regions in both datasets and 0.47 and 0.46 with CCF annotations, respectively. We also compared cell-type organization across independent cerebellar and isocortical spatial sections and found reproducible regional patterns across samples (**Supplementary Fig. 4b-c**). These findings indicate that the correspondence between inferred neighborhoods and tissue organization extends across independent sections and is not restricted to a single spatial sample.

Together, these analyses show that cellular neighborhoods derived from ProxiNet-predicted proximity relationships recover reproducible features of tissue architecture when spatial ground truth is available. This spatial validation provided an important basis for applying the same framework to dissociated scRNA-seq, where spatial coordinates are unavailable.

### Spatial calibration transfers cellular-neighborhood information to dissociated scRNA-seq

We next tested whether ProxiNet could transfer spatially learned cellular-proximity relationships to dissociated scRNA-seq^22^, where spatial coordinates are not available at inference. We applied the spatially calibrated model directly to scRNA-seq expression profiles and used the resulting pairwise proximity probabilities to construct a cellular-proximity graph. We then clustered this graph to identify inferred cellular neighborhoods (**Fig. 4a, Methods**). This analysis allowed us to test whether spatially learned relationships could be transferred beyond the spatial datasets used for model training.

**Figure 4.**
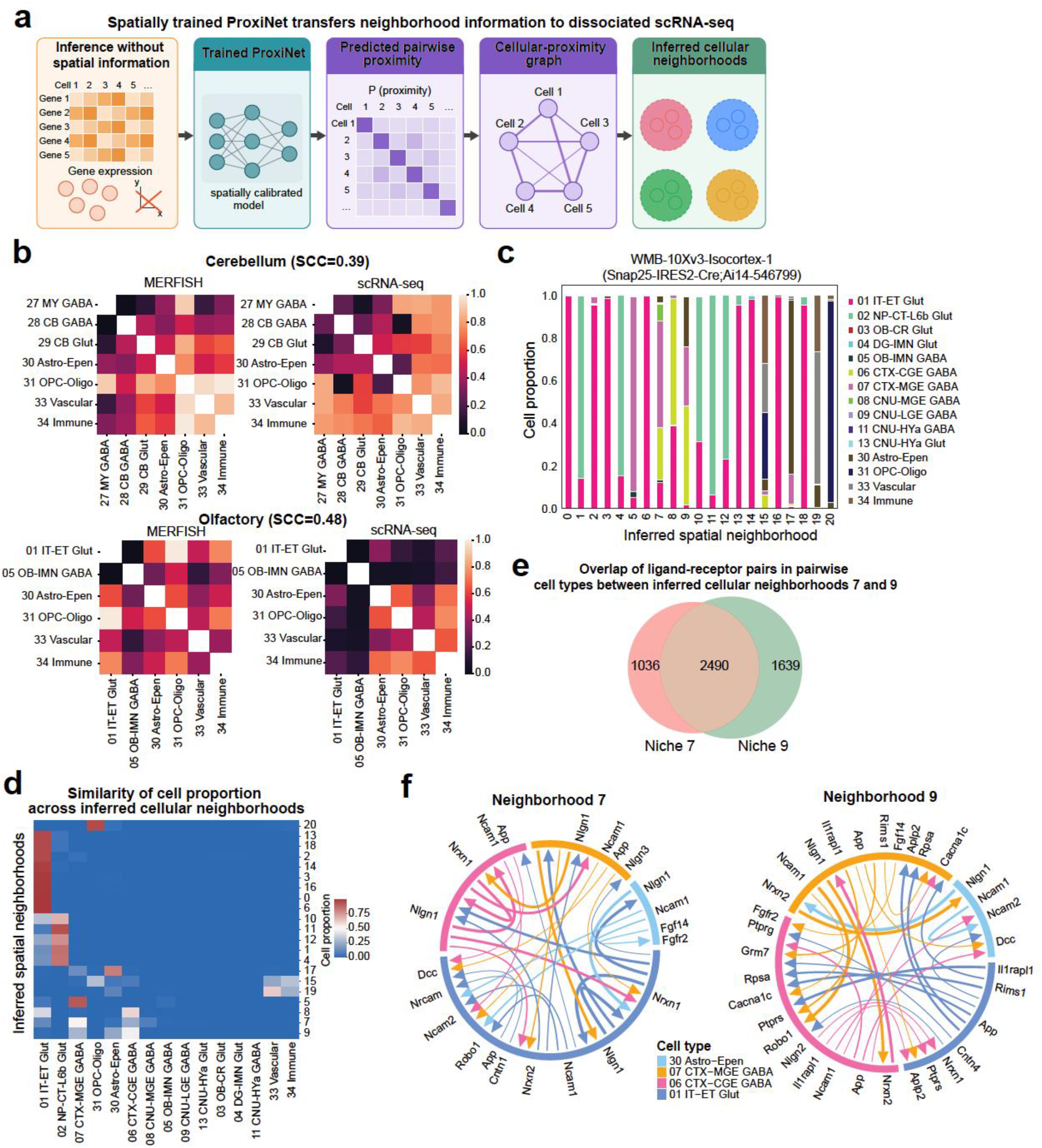
Spatial calibration transfers cellular-neighborhood information to dissociated scRNA-seq. **a**, Transfer of a spatially trained ProxiNet model to dissociated scRNA-seq. Gene-expression profiles from dissociated cells, without spatial coordinates or spatial information at inference, are processed by the spatially calibrated model to obtain pairwise proximity probabilities, construct a cellular-proximity graph, and identify inferred cellular neighborhoods. **b**, Concordance between cell-type proximity relationships measured in MERFISH and those inferred by ProxiNet from dissociated scRNA-seq for Cerebellum (SCC = 0.39) and Olfactory regions (SCC = 0.48). Heat maps show pairwise relationships among the indicated neuronal and non-neuronal cell classes; SCC is Spearman correlation coefficient. **c**, Cell-type composition of inferred cellular neighborhoods in the WMB-10Xv3-Isocortex-1 scRNA-seq sample (Snap25-IRES2-Cre;Ai14-546799). Stacked bars show the proportion of constituent cell types in each neighborhood. **d**, PCC-based comparison of cell-type composition across inferred Isocortex neighborhoods. **e**, Overlap of candidate ligand–receptor pairs across cell type pairs identified in inferred neighborhoods 7 and 9. Of the prioritized pairs (cell type 1-ligand-receptor-cell type2), 2,490 were shared between the two neighborhoods, whereas 1,036 and 1,639 were specific to neighborhoods 7 and 9, respectively. **f**, Candidate ligand–receptor interaction networks for neighborhoods 7 and 9, illustrating distinct communication programs involving Astro-Epen, CTX-MGE GABA, CTX-CGE GABA, and IT–ET Glut populations. Edge and arrow properties indicate the average expression level of ligand-receptor pairs and their directions between cell types. The color of the edges is defined by the ligand expressed cell types.

We first asked whether ProxiNet recovered cell-type proximity relationships that were consistent with those measured directly in spatial data. We compared cell-type proximity matrices derived from MERFISH with those inferred from dissociated scRNA-seq for matched brain regions. We found a Spearman correlation coefficient of 0.39 in Cerebellum and 0.48 in Olfactory, indicating that ProxiNet partially recovered the relative proximity structure among major neuronal and non-neuronal cell classes from dissociated transcriptomes alone (**Fig. 4b**). We observed similar concordance across additional brain regions, including Cortical Subplate, Pons, Pallidum, Medulla, Striatum, and Hypothalamus, with SCC values ranging from 0.23 to 0.37 (**Supplementary Fig. 5**). These results indicate that spatially calibrated proximity relationships transfer across anatomically distinct brain regions, although the degree of agreement varies across tissues.

We next examined whether the transferred pairwise relationships could resolve heterogeneous cellular neighborhoods within dissociated scRNA-seq. In the WMB-10Xv3-Isocortex-1 dataset, ProxiNet identified 21 inferred cellular neighborhoods with distinct compositions of excitatory neurons, inhibitory neurons, astrocytes, oligodendrocyte-lineage cells, vascular cells, and immune cells (**Fig. 4c**). We found that some neighborhoods were dominated by related cell populations, whereas others contained more heterogeneous combinations of neuronal and non-neuronal cell classes. We then compared neighborhood composition using pairwise similarity of cell-type proportions and found structured relationships among the inferred neighborhoods (**Fig. 4d**), suggesting that ProxiNet captures a continuum of related but compositionally distinct multicellular states rather than producing arbitrary graph partitions.

We further asked whether these neighborhood differences remained apparent at finer cell-subclass resolution. In the Isocortex dataset, we found that individual neighborhoods contained distinct combinations of transcriptionally defined excitatory, inhibitory, glial, vascular, and immune subclasses (**Supplementary Fig. 6a-b**). We observed a similar pattern in the WMB-10Xv3-STR dataset, where inferred neighborhoods contained different mixtures of striatal GABAergic populations, glutamatergic cells, astrocyte subtypes, oligodendrocyte-lineage cells, vascular populations, and immune cells (**Supplementary Fig. 6c-d**). These findings indicate that the inferred neighborhoods are not explained solely by broad cell-type identity and instead capture higher-order differences in multicellular composition but the shared spatial proximity across dissociated scRNA-seq datasets.

We next tested whether neighborhoods with related cellular composition could also differ in their associated candidate communication programs. We focused on Isocortex neighborhoods 7 and 9 and compared cell type-ligand-receptor-cell type tuples identified within each neighborhood. We found that 2,490 candidate tuples were shared between the two neighborhoods, whereas 1,036 were specific to neighborhood 7 and 1,639 were specific to neighborhood 9 (**Fig. 4e, Supplementary Table 6-7**). We then examined the corresponding ligand-receptor networks and found distinct interaction patterns involving Astro-Epen, CTX-MGE GABA, CTX-CGE GABA, and IT-ET Glut populations (**Fig. 4f**). These results suggest that neighborhoods with overlapping cellular composition can still differ in their associated candidate communication programs. Together, these analyses show that spatial calibration enables ProxiNet to transfer cellular-proximity information to dissociated scRNA-seq, recover heterogeneous cellular neighborhoods, and nominate neighborhood-associated molecular interactions without requiring spatial coordinates at inference.

### ProxiNet reveals age-dependent Alzheimer’s-associated remodeling of inferred cellular neighborhoods

We next asked whether ProxiNet could identify disease-associated changes in cellular-neighborhood organization from dissociated scRNA-seq. We applied the spatially calibrated model to a published entorhinal cortex dataset containing wild-type (WT) and Alzheimer’s disease (AD) samples at 2 and 8 months of age^23^. Major glutamatergic, GABAergic, astrocyte, oligodendrocyte, OPC, microglial, and endothelial populations were represented at both ages and in both conditions (**Fig. 5a**). We inferred pairwise cellular proximity independently within each age and condition and then clustered the resulting proximity graphs to define cellular neighborhoods. We found multiple neighborhoods with distinct cell-type compositions in each group (**Fig. 5b**), providing a framework to compare multicellular organization across disease progression.

**Figure 5.**
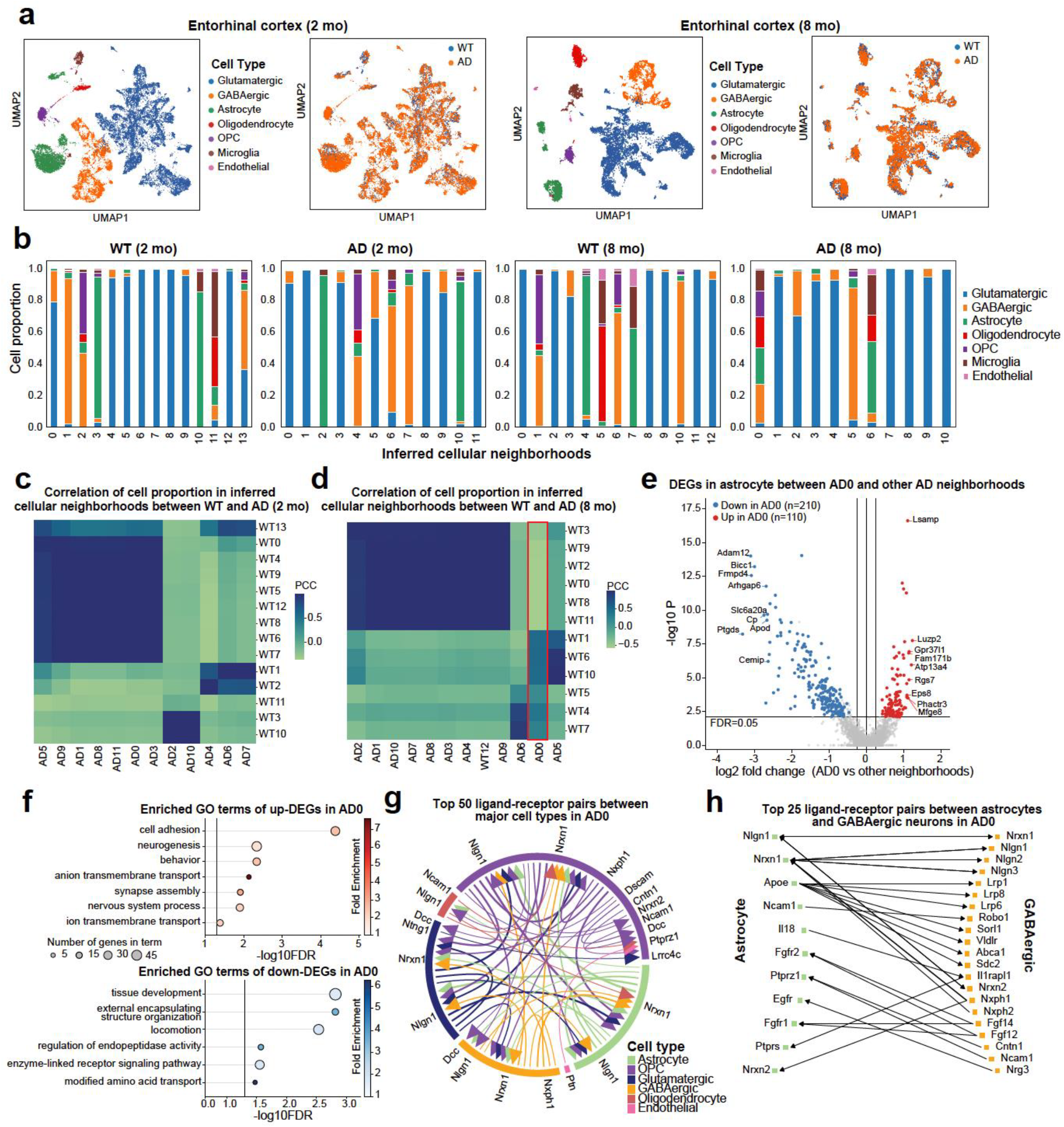
ProxiNet reveals age-dependent Alzheimer’s-associated remodeling of inferred cellular neighborhoods. **a**, Single-cell transcriptomic profiles of entorhinal cortex at 2 and 8 months from a published dataset. UMAP representations are colored by major cell type and by experimental condition (WT or AD) for each age. Major populations include glutamatergic neurons, GABAergic neurons, astrocytes, oligodendrocytes, OPCs, microglia, and endothelial cells. **b**, Cell-type composition of inferred cellular neighborhoods in 2-month WT, 2-month AD, 8-month WT, and 8-month AD samples. Each stacked bar represents the cellular composition of one inferred neighborhood. **c**, Correspondence between WT and AD neighborhoods at 2 months based on correlation of neighborhood cell-type composition. **d**, Correspondence between WT and AD neighborhoods at 8 months. AD neighborhood 0 (AD0; highlighted) shows reduced correspondence to WT neighborhoods relative to matched neighborhood structures. **e**, Differential gene-expression analysis of astrocytes in AD0 versus astrocytes from other AD neighborhoods at 8 months. The volcano plot shows log2 fold change and −log10P; 110 genes were upregulated and 210 genes were downregulated in AD0 at FDR < 0.05. Selected genes are labeled. **f**, Gene Ontology enrichment for genes upregulated (top) and downregulated (bottom) in AD0 astrocytes. Dot position denotes −log10FDR, dot color denotes fold enrichment, and dot size represents the number of genes assigned to each term. **g**, Top 50 candidate ligand–receptor pairs connecting major cell types within AD0 at 8 months. Colors denote the indicated cell classes; rank is based on the significance from LIANA package. **h**, Top 25 candidate ligand–receptor pairs between astrocytes and GABAergic neurons within AD0, highlighting candidate interactions including *Apoe*- and *Il18*-associated relationships alongside neuronal adhesion and synaptic interaction genes. Pairs were ranked by statistical significance.

We first examined the underlying cell-type proximity relationships across the four groups. ProxiNet predicted distinct patterns of pairwise proximity among major neuronal and non-neuronal populations in 2-month WT, 2-month AD, 8-month WT, and 8-month AD samples (**Supplementary Fig. 7a**). We then compared the cell-type composition of inferred neighborhoods between WT and AD. At 2 months, we found that most AD neighborhoods showed substantial compositional similarity to one or more WT neighborhoods (**Fig. 5c**). The hierarchical analysis of neighborhood composition showed a similar pattern, with WT and AD neighborhoods frequently grouping together according to their cellular composition (**Supplementary Fig. 7b**). These findings suggest that overall neighborhood organization remains broadly comparable between WT and AD at this earlier stage.

The pattern changed at 8 months. Although several WT and AD neighborhoods retained compositionally similar counterparts, we found that AD neighborhood 0, which we refer to as AD0, showed reduced correspondence to WT neighborhoods (**Fig. 5d**). The hierarchical analysis of 8-month neighborhood compositions also separated AD0 from many of the WT neighborhoods (**Supplementary Fig. 7c**). These results suggest that disease progression may be associated with remodeling of inferred cellular-neighborhood organization and nominate AD0 as an AD-associated multicellular state that becomes apparent at the later time point.

We next asked whether cells within AD0 exhibited transcriptional states that distinguished them from the same cell type in other AD neighborhoods. We first focused on astrocytes because they contributed prominently to the cellular composition of AD0. Differential expression analysis comparing AD0 astrocytes with astrocytes from other 8-month AD neighborhoods identified 110 upregulated and 210 downregulated genes at FDR < 0.05 (**Fig. 5e, Methods**). Among the differentially expressed genes were *Lsamp, Luzp2, Gpr37l1, Mfge8, Cp, Ptgds, Apod*, and *Cemip*. Several of these genes have previously been linked to astrocyte heterogeneity or AD-associated astrocyte functions. In particular, *Mfge8* is expressed by astrocytes and has been implicated in astrocyte-mediated regulation of synapse elimination in AD, while *Apod, Cp*, and *Gpr37l1* have been associated with astrocyte lipid metabolism, metal homeostasis, or glutamate and neuronal signaling in neurodegenerative contexts^24,25^. Gene Ontology analysis showed that genes upregulated in AD0 astrocytes were enriched for ion transmembrane transport, nervous system processes, synapse assembly, neurogenesis, and cell adhesion, whereas downregulated genes were associated with modified amino acid transport, receptor signaling, regulation of endopeptidase activity, extracellular structure organization, and tissue development (**Fig. 5f**). These functions are consistent with established roles of astrocytes in ion and neurotransmitter homeostasis, synapse formation and elimination, and cell-cell interactions, all of which are altered in AD^26,27^. We therefore found that astrocytes within the AD0 neighborhood occupy a distinct transcriptional state, suggesting that cellular state in AD is associated with the inferred multicellular context in which the cells reside.

We next examined candidate intercellular communication programs associated with AD0. Using LIANA, we ranked ligand-receptor relationships among the major cell populations within AD0 and visualized the top 50 candidate interactions (**Fig. 5g, Supplementary Table 8**). We found an extensive network connecting astrocytes, OPCs, glutamatergic neurons, GABAergic neurons, oligodendrocytes, and endothelial cells. We then focused on interactions between astrocytes and GABAergic neurons and found candidate relationships involving genes such as *Apoe, Il18, Nrxn1, Nlgn1, Ptprz1*, and several neuronal adhesion and signaling molecules (**Fig. 5h**). These results nominate candidate communication programs associated with the AD0 neighborhood, including relationships between astrocytes and neuronal populations. Because these analyses are based on ligand-receptor expression and inferred cellular proximity, they identify candidate interactions rather than demonstrating active signaling between the corresponding cells.

Finally, we asked whether the transcriptional changes associated with AD0 extended beyond astrocytes. We compared GABAergic neurons in AD0 with GABAergic neurons from other 8-month AD neighborhoods and identified 91 upregulated and 150 downregulated genes (**Supplementary Fig. 7d**). The upregulated genes were enriched for processes including modulation of chemical synaptic transmission, synapse assembly, ion transport, calcium-mediated signaling, and cell adhesion, whereas the downregulated genes were enriched for synapse organization, neuronal projection morphogenesis, membrane potential regulation, neuronal migration, and cell migration (**Supplementary Fig. 7e**). We therefore found distinct transcriptional programs in more than one cellular population within AD0, consistent with AD0 representing a multicellular neighborhood state rather than a transcriptional change restricted to astrocytes. Together, these analyses show that ProxiNet can identify age-associated remodeling of inferred cellular neighborhoods and nominate molecular programs associated with disease-specific multicellular organization from dissociated single-cell transcriptomes.

## Discussion

We developed ProxiNet to transfer cellular-proximity information learned from spatial transcriptomic references to dissociated single-cell transcriptomes. Rather than treating coordinate reconstruction as the primary objective, ProxiNet formulates tissue organization as a supervised pairwise prediction problem. We found that proximity prediction generalized across held-out brain regions and across MERFISH, STARmap, and Slide-seq. We also found substantial zero-shot transfer between spatial technologies, while target-platform fine-tuning further improved performance. These results suggest that ProxiNet captures proximity-associated transcriptomic features that extend beyond individual datasets while retaining sensitivity to platform-specific differences. We further found that pairwise predictions could be assembled into cellular-proximity graphs that recovered anatomical organization in spatial datasets and heterogeneous neighborhoods in dissociated scRNA-seq.

ProxiNet complements methods that reconstruct absolute spatial position. Tangram and CytoSPACE align dissociated cells to measured spatial references, while novoSpaRc and CeLEry infer spatial locations from transcriptomic information^4–7^. These approaches are well suited when spatial coordinates or cell-to-location assignments are the desired output. ProxiNet instead predicts whether two cells are likely to have occupied nearby tissue environments and represents these predictions directly as weighted graph edges. Our auxiliary analyses showed that ProxiNet representations also retain information sufficient for coordinate recovery, suggesting that coordinate reconstruction and relational proximity inference capture overlapping but distinct aspects of spatial organization.

ProxiNet also differs from recent approaches that transfer spatial context through latent representations. CellContrast learns a contrastive embedding in which spatially proximal cells are similar, ENVI transfers covariance-based niche representations, and Nicheformer predicts spatial properties such as niche labels and neighborhood composition^8–10^. ProxiNet directly optimizes pairwise proximity probabilities, which provides an explicit relational output that can be used as an edge weight for downstream graph analysis. This formulation also allows us to trace individual proximity predictions back to transcriptomic features. We found that highly attributed genes were enriched for ligand-receptor relationships and that spatially attributed genes nominated broader cell-type-specific transcriptional modules. These results suggest that proximity predictions capture biologically structured molecular programs rather than only broad cell-type identity. At the same time, model attribution identifies features associated with predictions rather than causal determinants, and ligand-receptor analyses nominate candidate interactions rather than demonstrating active signaling.

The neighborhood analyses extend this relational framework from individual cell pairs to higher-order tissue organization. We found that communities derived from ProxiNet proximity graphs formed spatially coherent domains and showed reproducible concordance with major brain regions and CCF annotations. The correspondence was substantial but incomplete, suggesting that inferred neighborhoods reflect anatomical organization without simply reproducing predefined region labels. This distinction is relevant because cellular neighborhoods can capture multicellular organization at a finer scale than conventional anatomical domains. Previous spatial studies have similarly shown that local cellular neighborhoods can distinguish biologically meaningful tissue states that are not captured by cell abundance alone^2^. We further found that inferred neighborhoods in dissociated Isocortex and Striatum contained distinct combinations of neuronal and non-neuronal subclasses and that compositionally related neighborhoods could differ in candidate communication programs. These findings suggest that inferred neighborhood context provides an additional level of organization beyond conventional cell-type annotation.

Our Alzheimer’s disease analysis illustrates how this framework can be used to study condition-associated remodeling from dissociated data. Single-cell studies have established that AD involves heterogeneous and cell-type-specific transcriptional responses, including altered neuronal, astrocyte, oligodendrocyte, and inflammatory programs^28,29^. Consistent with the importance of cellular context in disease progression, we found that WT and AD neighborhoods were broadly comparable at 2 months, whereas an AD-associated neighborhood, AD0, emerged more clearly at 8 months. Astrocytes in AD0 showed transcriptional differences from astrocytes in other AD neighborhoods, and GABAergic neurons showed additional neighborhood-associated changes. These results suggest that disease-associated transcriptional states can be organized within multicellular contexts rather than occurring uniformly across all cells of a given type. We also identified candidate ligand-receptor programs involving astrocytes, neurons, and other cell populations within AD0, providing hypotheses for subsequent spatial or experimental validation.

Several limitations define the current scope of ProxiNet. First, we trained and extensively evaluated the model in mouse brain datasets. Although we found generalization across anatomical regions and three spatial technologies, testing additional organs, species, and disease contexts will be necessary to establish broader transferability. Second, ProxiNet infers proximity from transcriptomic state and does not directly measure physical adjacency in dissociated samples. Predicted edges should therefore be interpreted as inferred relationships rather than evidence of direct cell-cell contact. Third, neighborhood assignments depend on proximity and graph-clustering parameters, and future work should further characterize their stability across parameter choices. Fourth, gradient attribution and ligand-receptor analyses identify associations and candidate communication programs rather than causal mechanisms. Finally, targeted spatial platforms measure only part of the transcriptome, so extending spatially attributed genes to whole-transcriptome modules depends on the correspondence between spatial and dissociated cell states.

Future work can address these limitations by training on increasingly diverse spatial atlases and explicitly modeling technology-specific variation to improve zero-shot transfer. Additional molecular and imaging modalities could also help distinguish cellular relationships that are difficult to resolve from RNA alone. More generally, the pairwise formulation provides a flexible interface between spatial atlas learning and graph-based analysis of dissociated cells. By treating cellular proximity as a transferable relational quantity, ProxiNet provides a framework for recovering multicellular organization from existing single-cell datasets and for prioritizing spatial relationships for experimental validation.

## Methods

### Preprocessing of datasets

#### MERFISH

We used the whole-mouse-brain MERFISH dataset from Zhang et al. (2023), which profiled 1,122 genes in approximately 10 million cells across the adult mouse brain^11^. The dataset includes more than 5,000 transcriptionally defined clusters organized into 34 major cell classes and 338 subclasses. We obtained the full expression matrix together with spatial coordinates and preannotated cell-type metadata.

For ProxiNet training and evaluation, we separated each MERFISH specimen into individual brain sections so that each sample contained cells from a single tissue section. This resulted in 239 samples, which we randomly divided into training, validation, and test sets using a 7:1:2 ratio (**Supplementary Table 1**). In total, 167 samples were used for training, 24 for validation, and 48 for testing. We obtained spatial coordinates and cell-type annotations directly from the accompanying metadata. For each cell pair, we calculated the spatial distance and assigned a binary proximity label using a distance threshold of 20 µm, with pairs below the threshold labeled as proximal and all other pairs labeled as non-proximal. Unless otherwise specified, all model evaluation and downstream analyses were performed using test samples that were not used during model training or validation.

#### STARmap

We used the whole-mouse-brain and spinal-cord STARmap dataset from Shi et al. (2023), which profiled 1,022 genes in three-dimensional space across approximately 1.09 million cells and annotated 230 cell types and 106 anatomical regions^12^. Because the dataset contained fewer tissue sections than the MERFISH dataset, we divided the available sections into 13 training samples, 2 validation samples, and 5 test samples (**Supplementary Table 2**).

For consistency with the other spatial datasets, we used the first two spatial dimensions, x and y, for proximity - label generation. We extracted these coordinates directly from the metadata and calculated pairwise spatial distances within each sample. As for MERFISH, we assigned binary proximity labels using a distance threshold of 20 µm.

#### Slide-seq

We used the mouse whole-brain Slide-seq dataset from Langlieb et al. (2023), which combines spatial transcriptomic profiling with high-throughput single-nucleus RNA sequencing^13^. The dataset contains 4,388,420 nuclei, with a median of 4,884 unique molecular identifiers per nucleus and 1,937 transcriptionally defined clusters. We prepared 69 samples for training, 11 for validation, and 21 for testing (**Supplementary Table 3**).

Because Slide-seq provides transcriptome-wide measurements and contains 29,517 genes, we restricted the input gene set to genes that were also measured in MERFISH or STARmap to facilitate model training and cross-technology comparison. We calculated pairwise spatial distances within each sample and used a distance threshold of 10 µm to define positive proximity labels.

#### Whole-mouse-brain scRNA-seq

We used the WMB-10Xv3 scRNA-seq dataset from Yao et al. (2023), which profiled approximately 7 million cells, with more than 2.5 million cells retained after quality-control filtering^22^. The dataset contains measurements for 32,285 genes, including 8,460 genes identified as differentially expressed across the atlas. Importantly, the WMB-10Xv3 atlas uses the same class and subclass annotation framework as the corresponding MERFISH whole-brain atlas, enabling consistent comparison between dissociated and spatial datasets.

The dataset was organized into 12 major brain regions: Olfactory, Cortical Subplate, Hippocampus, Isocortex, Striatum, Pallidum, Hypothalamus, Thalamus, Midbrain, Pons, Medulla, and Cerebellum. We treated cells from each brain region as a separate sample for graph construction. Raw expression counts were used to construct the heterogeneous cell-gene graphs and to generate cell representations using the MERFISH-trained ProxiNet model. No spatial coordinates were provided to ProxiNet during inference. Pre-normalized expression profiles were used only for downstream analyses, including UMAP visualization and ligand-receptor analysis.

#### Alzheimer’s disease scRNA-seq

We used the entorhinal cortex scRNA-seq datasets at 2 and 8 months of age from Bartas et al. (2025) ^23^. The dataset contains eight samples spanning wild-type (WT) and Alzheimer’s disease (AD) conditions at the two ages. The 2-month dataset contains 8,435 WT cells and 13,659 AD cells, whereas the 8-month dataset contains 6,069 WT cells and 5,011 AD cells. Preannotated major cell types included glutamatergic neurons, GABAergic neurons, astrocytes, oligodendrocytes, oligodendrocyte precursor cells (OPCs), microglia, and endothelial cells.

We used raw gene-expression counts to construct cell-gene graphs and applied the MERFISH-trained ProxiNet model to generate cell representations and infer pairwise cellular proximity within each age and condition. We then constructed proximity graphs and inferred cellular neighborhoods separately for 2-month WT, 2-month AD, 8-month WT, and 8-month AD cells. Pre-normalized expression profiles were used for downstream analyses.

### Construction of the cell-gene bidirected graph

We aimed to mimic the cell-cell interaction network with a bidirected graph, where information (molecular signals) can flow in both directions from one cell to another. For each single cell measured in the spatial transcriptomics sample (M cells and N genes measured), we initialized a cell node *c*_*i*_ representing the cell itself as well as a feature vector 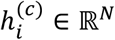 with its gene expression profile. For each gene measured, we created a gene node *g*_*j*_ for its representing gene with a feature vector 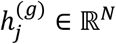 by one-hot encoding. To simulate the process of cell-cell communication, an edge with weight *w*_*ij*_ was created between *c*_*i*_ and *g*_*j*_ if gene *j* has expression level *w*_*ij*_ in cell *i*. Hence, the information will first flow from *c*_*i*_ to *g*_*j*_, indicating the participation of gene *j* in sender activities, and then flow from *g*_*j*_ to another cell node *c*_*k*_, indicating gene *j* operates in receiver activities. With the set of full cell and gene node *V* = {*c*_1_, *c*_2_, …, *c*_*M*_, *g*_1_, *g*_2_, …, *g*_*N*_} and the set of all possible edges *E* = {(*c*_*i*_, *g*_*j*_) | *g*_*j*_ *is expressed in c*_*i*_} constructed, the final bidirected graph *G* = (*V, E*) was formed which simulates a complex and multi-layer cell-cell interaction network, where cell signals can transverse from one cell to another cell through all possible cell-gene-cell routing combinations. To make fair evaluation between different datasets, we set a common gene list containing 1,712 genes (**Supplementary Table 4**) which is the union of all genes measured in MERFISH and STARMap, resulting in identical gene nodes between graphs from different sources and making the trained models more generalizable. For each graph, the edge weight *w*_*ij*_ is first normalized with target sum 1e4 and then log-transformed.

Gene-gene interaction also exists and the expression of one gene may affect another gene-by-gene regulation and ligand-receptor interaction. We also provided an option to create edges by biological priors with unit weight between gene node *g*_*j*_ and *g*_*k*_ if (i) gene *j* is a transcription factor of gene *k* or vice versa; (ii) they are involved within the same signaling pathway; (iii) they act as a ligand-receptor pair. With created gene-gene linkages, information may flow from one gene to another without going through cells. We obtained biological priors from public gene regulatory datasets including the ChEA Transcription Factor Targets^30^ and the KEGG PATHWAY Database^31^. For the analysis in this study, we used the standard graphs with cell-gene edges.

### Framework of ProxiNet

ProxiNet is a multi-step framework composed of multiple modules for different purposes. Once the bidirected - graph is constructed, a graph-neural network (GNN) module named ProxiGNN is used to project the input feature vector *h* ∈ ℝ^*N*^ into a low-dimensional feature vector *h* ∈ ℝ^256^ and eventually generate an embedding vector *z* for each cell node and gene node. Predictor and Embedder are separate multilayer perceptrons designed for further processing of the outputs of ProxiGNN. Predictor takes in aggregated embeddings from sampled pairs of cell nodes and outputs proximity values *p*. Embedder uses all embeddings as inputs but generates 2D coordinates (*x, y*) respectively. We trained a separate set of ProxiGNN and the following multilayer perceptron for each task to ensure the models achieve both accuracy and specificity.

#### ProxiGNN

ProxiGNN, consisting of several repeated units, acts as the core of ProxiNet. Each unit is formed by three consecutive HeteroGraph convolutional modules (HeteroGraphConv)^32^ *H*(*h*) linked by residual connection.

HeteroGraphConv, working on heterogeneous graphs with three types of edges: cell node to gene node, gene node to cell node and optional gene node to gene node. To properly aggregate information by edge categories, we created three separate channels with each one operating an independent GraphSAGE convolutional layer (SAGEConv)^33^. For the feature vector 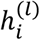 of node *i* at layer *l* (either cell node 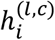 or gene node 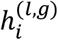 SAGEConv calculates the vector 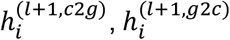 and 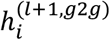 for the next layer *l* + 1 separately by the same operation:

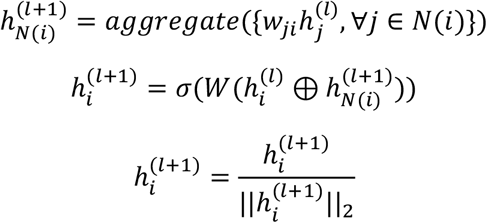

where *aggregate*(*h*) is the convolution function used to aggregate feature vectors in the neighborhood *N*(*i*) and is set as “mean” for the first HeteroGraphConv module and “max-pooling” for the remaining two modules. *w*_*ji*_ is the edge weight between node *j* and node *i. W* is the linear transformation mapping the aggregated feature vector back to a certain dimension. In the first SAGEConv layer, we reduce the dimension from 1,712 to 256, resulting in a feature vector with a lower dimension. For all following layers we keep the hidden feature vectors mapped back to the same hidden dimension. σ(*h*) denotes the ReLU activation function. A dropout rate of 0.25 is applied to each layer to prevent overfitting while normalization is used to prevent extreme values in the feature vectors. Eventually, we averaged the output of cell to gene channel 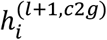, gene to cell channel 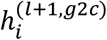 and gene to gene channel 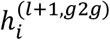 as the final feature vectors for cell node and gene node. Since incoming edges of cell nodes are from only gene nodes while incoming edges of gene nodes are possible from both genes and cells, the updated vectors are equal to:

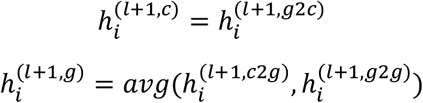

We iterated through all cell nodes and gene nodes and updated their feature vector after each SAGEConv layer. Before feeding them into the next HeteroGraphConv block, we performed a residual connection to avoid vanishing gradient by:

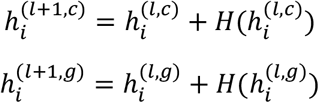

The unit of HeteroGraphConv modules are repeated three times before passing the final output vector — embedding *z* to the downstream modules (Predictor and Embedder) for specific tasks. In all, ProxiGNN processes input feature vectors with gene expression values and generates embeddings with low-dimension representations by: *h* ∈ ℝ^1712^ → *h* ∈ ℝ^256^ → *z* ∈ ℝ^256^ through graph convolution operations.

#### Predictor

Predictor *P*(*z*) is a standard multilayer perceptron with two linear layers. To predict proximity between a cell pair *i* and *j*, the input is a concatenated embedding vector with dimension 512. The first linear layer with ReLU activation and 0.25 dropout rate is used to reduce the dimension of the vector into 256, while the second linear layer with sigmoid activation directly produces the proximity. The final loss function is the binary cross entropy (BCE) between the ground truth label 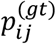 and the output value 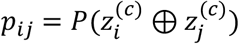:

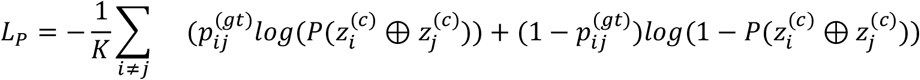

where 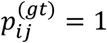 if cell *i* and cell *j* are in proximity otherwise 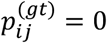. *K* is the number of cell pairs evaluated in each batch.

#### Embedder

Embedder *E*(*z*) is also a standard multilayer perceptron with different numbers of layers and dimensions. It uses 5 linear layers to map input embedding vectors into hidden vectors and the output vector with dimension 128, 64, 32, 16 and 2. The output vector 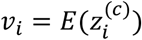 with 2 entries (*x*_*i*_, *y*_*i*_)^*T*^ can be viewed as predicted coordinates and is comparable with ground truth coordinates 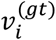. The loss function is defined as the mean squared error (MSE) between the pairwise distance matrices from both the ground truth and the prediction, which follows the same criteria defined by previous studies^18^:

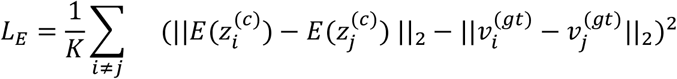

where *K* is the number of cell pairs evaluated in each batch.

#### Optimizer and regularization

We used the ADAM optimizer^34^ with initial learning rate 1e-4. The batch size is set to be 1,024 for all cases while early stopping is enabled when the average loss of validation set is not dropping after a certain number of epochs (for example 10 to 50 based on different tasks and datasets). We also applied further training with a smaller learning rate equals 1e-5 to refine the trained models. All models generally converge within 1,000 epochs. For Predictor training, each training batch contains the same number of positive cell pairs (cell pairs that are in proximity) and negative cell pairs (cell pairs that are not in proximity), both categories were randomly sampled from a precalculated cell pair list. Here we separately trained models on positive cell pairs can be from all possible cell pairs in proximity and only proximal cell pairs belonging to different subclasses. The first model was used for proximity-related evaluation and the second model was used for other downstream analysis like mapping between samples and niche detection. For Embedding training, we randomly segmented and sampled 1,024 cells together with their expression vectors from the training datasets for each training batch.

To avoid overfitting, we applied both L1 and L2 regularization during training to. L1 regularization adds a penalty term to the final loss function proportional to the sum of absolute values of all coefficients. Using ProxiGNN and Predictor as examples, the final loss is added by additional terms:

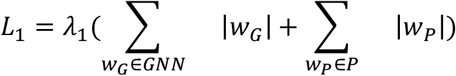

where *w*_*G*_ is any coefficient belonging to the ProxiGNN module *GNN* and *w*_*P*_ is any coefficient belonging to Predictor *P*. L2 regularization adds a similar penalty but with squared values instead of absolute values by:

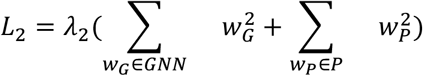

The final aggregated loss is:

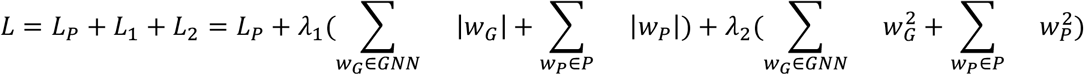

where both λ_1_ and λ_2_ are set to be 1e-5. Both regularization methods prevent large parameters and enhance the generalizability of the models. In Embedder training we replaced *w*_*P*_ in the formula by *w*_*E*_ to calculate loss terms *L*_*E*_. All modules in the ProxiNet framework were implemented by PyTorch with CUDA extension^35^ and deep graph library (DGL)^32^. All models with different setups are generally available for training using one NVIDIA-A100 GPU with memory no larger than 80GB.

### Benchmarking methods

#### Logistic Regression (LR)

Logistic regression (LR) is a linear classification method learned on linear combination of input variables. In our case, each variable indicates the expression level of a gene which contributes to the final probability output through factor multiplication and sigmoid conversion. A 0.5 threshold is then set to turn the probability into categories 0 and 1. The linear factors as parameters were optimized on the same training samples and then used for evaluation on the same testing samples.

#### Support Vector Machine (SVM)

Support vector machine (SVM) is a classification method aiming to find a linear decision boundary that divides data points into binary categories in the hyperspace^16^. Unlike logistic regression which uses a maximum likelihood estimation to learn the parameters, SVM trains the classifier solely with the closest points from both classes. Here we treated the concatenated and normalized gene expression vectors of each cell pair as input and their corresponding proximity as output. The hard-margin was used to separate points belonging to positive pairs and points belonging to negative pairs.

#### Multilayer Perceptron (MLP)

Multilayer Perceptron (MLP) is a widely-used classifier made up of fully connected layers for non-linear classification problems^15^. Here we adopted a standard two-tower MLP framework, where each tower consists of three fully-connected layers with 4, 16 and 64 scaling ratios and ReLU activations. Here we used the normalized expression vectors of each cell pair as input for both towers and the final output is calculated using the cosine similarity between their output vectors. The model was trained on the same set of training samples and was later evaluated on the same set of testing samples.

#### Convolutional Neural Network (CNN)

Convolutional Neural Network (CNN) facilitates convolutional operation by kernels during feature extraction and has been proved an efficient approach for non-linear classification^14^. We implemented CNN using the same framework as MLP where each of the two tower processes normalized gene expression and the final scores are calculated by cosine similarity. We used two intermediate 1D convolutional layers with channels equals 4, 16 and an output 1D convolutional layer whose stride equals 2 and kernel size equals 3.

#### Variational Autoencoder (VAE)

Variational Autoencoder (VAE) is a neural network architecture using variational Bayesian methods to map data points into a latent distribution^17^. For the encoder and the decoder, we used the same parameter setting as MLP where three dense layers with 4, 16 and 64 scaling ratios embed input vectors, the normalized expression profiles concatenated from both cells, into latent mean and latent standard deviation before they are reconstructed to the original sizes. The latent mean vectors are then feed-forwarded into a downstream classifier which compresses the vectors by 4 and further to the final output with sigmoid transformation. The binary cross entropy is used for loss calculation between the output values and labels.

#### Tangram

Tangram^5^ learns a probabilistic mapping of single-cell transcriptomes to spatial reference locations by matching shared gene expression and cell-density distributions. We used Tangram to map each query sample to its reference pair (**Supplementary Table 5**) and the full 1,122 genes in the MERFISH dataset. The mapped coordinates were later reconstructed by assigning query cells to its best mapped cells in the reference. The package is available at https://github.com/broadinstitute/Tangram.

#### NovoSparRc

NovoSpaRc^36^ uses optimal transport to reconstruct cell locations under the assumption that nearby cells tend to have similar gene-expression profiles, with optional spatial marker-gene information. We used the same reference-query mapping table (**Supplementary Table 5**) and created an atlas matrix with the top 100 highly variable genes out of the full 1,122 genes for each reference. The reconstruction was later run with an alpha_linear of 0.8, an epsilon of 5e-3 and a number of neighbors equals 30 for both source and target. The package is available at https://github.com/rajewsky-lab/novosparc.

#### CytoSPACE

CytoSPACE^7^ assigns individual single-cell transcriptomes to spatial transcriptomic locations by solving a constrained linear assignment problem with a cost based on gene-expression correlations. Since the mapping of CytoSPACE requires the same class categories in both query and reference. We first filtered each query and reference pair from the reference-query mapping table (**Supplementary Table 5**) to ensure consistency. The coordinates were then mapped using the single-cell resolution mode with a number of selected spots equal 10,000. The package is available at https://github.com/digitalcytometry/cytospace.

#### CeLEry

CeLEry^6^ predicts cell coordinates from gene expression using a supervised neural network trained on spatial transcriptomics data with known locations. To make fair comparisons with ProxiNet, we also trained a single CeLEry model on all 167 MERFISH training samples and evaluated it on the 48 MERFISH testing samples (**Supplementary Table 1**). We used the original setting for the parameters with a batch size of 64 and a maximum number of epochs equals 3,000 to fit a larger training set. The package is available at https://github.com/QihuangZhang/CeLEry.

#### CellContrast

CellContrast^8^ learns expression representations from spatial transcriptomics data through contrastive training, so that similarity between representations reflects spatial proximity among query cells. Due to the memory constraints, we trained three separate CellContrast models for individual MERFISH donors (one for Zhuang-ABCA-1, one for Zhuang-ABCA-2 and one for Zhuang-ABCA-3 and Zhuang-ABCA-4) (**Supplementary Table 1**). All models were trained with the original parameters with a batch size of 64 and a maximum number of epochs equals 10,000. The package is available at https://github.com/HKU-BAL/CellContrast.

#### LUNA

LUNA^18^ is a diffusion-based model to infer cell coordinates without templates. We used LUNA to infer two-dimensional cell coordinates from gene-expression profiles, following the authors’ implementation with default model hyperparameters. The model used eight transformer layers, 16 attention heads and 1,000 diffusion steps with a cosine noise schedule. We adapted the input to the 1,122 gene-expression features in the MERFISH dataset and trained for 1,000 epochs with a reduced training batch size from six slices to one slice to accommodate GPU memory constraints, a learning rate of 5e-4 and weight decay of 1e-12 (**Supplementary Table 1**). Validation was disabled, and checkpoints were saved every 250 epochs. At inference, LUNA generates coordinates through iterative denoising conditioned on expression profiles, without requiring a spatial reference slice. The package is available at https://github.com/mlbio-epfl/luna.

### Evaluation metrics

#### Accuracy

Accuracy (ACC) is used to evaluate classification performance. In proximity prediction, the Predictor output of cell pair *i* and *j*, in other words 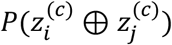, was first converted to a binary label with a 0.5 threshold:

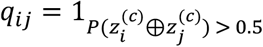

where 1 is the indicator function. Then we compared every ground truth pair *p*_*ij*_ against the predicted value *q*_*ij*_ and obtain the numbers of true positive (TP), false positive (FP), true negative (TN) and false negative (FN). The binary accuracy is quantified as:

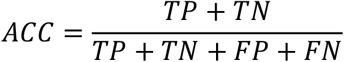

#### Area Under the Precision-Recall Curve (AUPR)

Area under the precision-recall curve (AUPR) is defined as the area bounded by the precision-recall curve and two axes. The precision-recall curve is formed by calculating the precision against recall at different thresholds.

For a given threshold *t*, we first converted the Predictor output to binary labels by 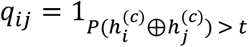. After that, TP, FP, TN, FN could be directly found by comparing ground truth *p*_*ij*_ and prediction *q*_*ij*_. The precision and recall is defined as:

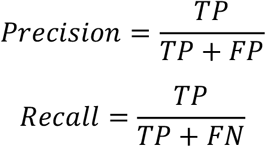

We calculated both values for every *t* ranging from 0 to 1 and the final curve was constructed. The area under the curve is sequentially denoted as AUPR. A random classifier has a 0.5 AUPR while a good classifier has an AUPR close to 1.

#### Area Under the Receiver Operating Characteristic (AUROC)

Similar to AUPR, Area under the receiver operating characteristic is defined as the area bounded by the receiver operating characteristic curve and two axes. This curve is formed by calculating the true positive rate (TPR) against the false positive rate (FPR) at different thresholds. After obtaining TP, FP, TN, FN from *p*_*ij*_ and *q*_*ij*_, we calculated TPR and FPR by:

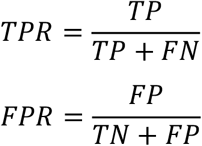

We evaluated both values across every threshold and the final curve was constructed. AUROC is then formed by the area under the curve. A random classifier has a 0.5 AUROC while a good classifier has an AUROC close to 1.

**Jensen-Shannon Distance (JSD)**

Jensen-Shannon Distance (JSD) is used to quantify whether two probability distributions are similar or not and has been applied to evaluate the local environment of cells^8,37^. In our case, the distributions are discrete distributions of cell types. For cell *i* from a sample with M cells, we calculated its 200 nearest neighbors and count their cell type distributions from both ground truth and predicted coordinates, labeled as 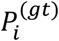 and 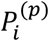. JSD is defined as:

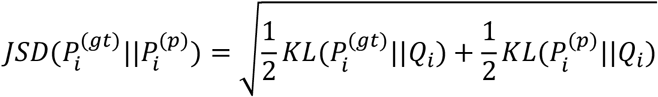

where *KL*(*X*||*Y*) is the Kullback-Leibler divergence between distribution *X* and *Y* while *Q*_*i*_ is the average distribution of 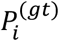 and 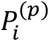. We averaged JSD values from all cells to obtain the JSD for the whole sample:

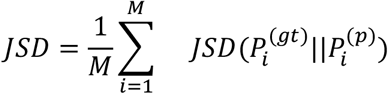

A smaller JSD indicates a better locally alignment between the ground truth and the prediction.

#### Spearman’s Rank Correlation Coefficient (SCC)

Spearman’s Rank Correlation Coefficient (SCC) is a non-linear correlation measurement used by previous study^8,18^ to evaluate reconstructed coordinates. In our case, we aimed to compare how pairwise distances are similar between the ground truth and prediction. For cell *i* from a sample with M cells, we calculated its spatial distance with all cells in the sample, resulting in a distance vector *d*_*i*_ with dimension M. The ground truth vector 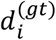 and the prediction vector 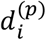 were then converted to ranks 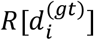 and 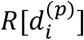 to calculate their Pearson correlation coefficient:

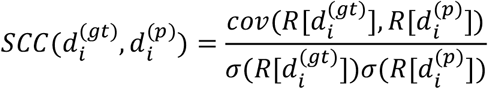

where *cov*(*x, y*) and σ(*x*) denotes covariance and standard deviation functions with vectors *x* and *y* as inputs. The SCC for the whole sample was then similarly calculated by:

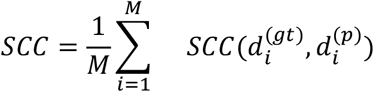

A larger SCC means there is a better global alignment between the ground truth and the prediction.

#### Normalized Mutual Information (NMI)

Normalized Mutual Information (NMI) measures how similar two groups of clusters are by evaluating their mutual information over the average of their Shannon entropy values^38^. Assume the joint and marginal probability mass functions of two distribution *X* and *Y* are *p*(*x, y*), *p*(*x*) and *p*(*y*), the mutual information is calculated by:

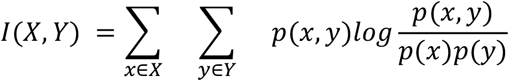

With Shannon Entropy for *X* and *Y* calculated by *H*(*X*) = − ∑_*x*∈*X*_ *p*(*x*)*logp*(*x*) and *H*(*Y*) = − ∑_*y*∈*Y*_ *p*(*y*)*logp*(*y*), the final NMI is formulated by:

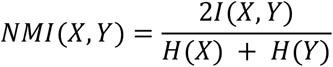

A higher NMI score indicates the two groups share more mutual information and have more similar clustering assignments.

### Gradient analysis

We used gradient analysis to quantify the contribution of graph nodes^19^. In our model with only cell-gene edges, ProxiGNN first aggregates information from every cell node’s direct gene node neighbors to generate an embedding vector *z* = *G*(*h*) which is then feedforwarded to Predictor to generate output *p* = *P*(*z*). The accumulated gradient of a node can be calculated by backpropagation from the output to the node itself using the chain rule with the final value *p* and the message passing function *p* = *P*(*G*(*h*)). For cell *i* with feature vector 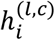 at layer *l* in ProxiGNN specifically, its gradient is calculated by summing all gradients coming from its previous layer together with its neighboring genes *N*(*i*):

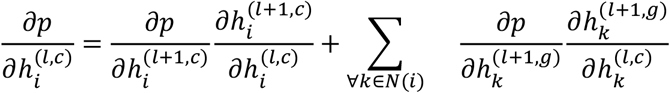

Similarly for gene *j* with feature vector 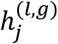 at layer, we have:

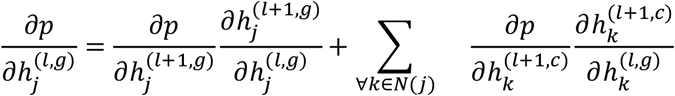

Since each cell node is only connected to gene nodes, the gradients at the incoming edges to the cell nodes measure how each gene affects the proximity between the cell pair. Respectively we may calculate the partial derivatives accumulated to the initial layer for all gene nodes’ representation: 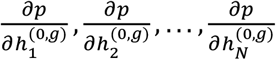. These gradients measure how significantly these genes are contributing to the final proximity. A large 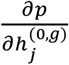 means gene *j* plays an important role in forming the cell interaction pair. In general, the top 50 genes for each cell type are defined as proximity-driven genes and kept for downstream analysis.

To judge whether detected proximity-driven genes are meaningful, we used Fisher’s exact test to test whether these genes are associated with ligand-receptor pairing genes from the mouse consensus database. Using full protein-coding genes as the background set of genes, we built a contingency table by counting the numbers of genes in proximity-driven genes and the database, either or not. A p-value < 0.05 indicates the cell pair has a proximity-driven gene set significantly associated with potential ligand-receptor interaction. For each MERFISH testing sample, we applied the test on all possible cell type pairs presented not less than 100 times.

### Gene co-expression module identification

We identified gene co-expression modules separately within cell populations using the Allen Institute whole-mouse-brain scRNA-seq atlas^22^, restricting the analysis to WMB-10Xv3. We used the precomputed log2(CPM+1) expression values and analyzed cells at two grouping levels: Allen subclass and class by anatomical division. Groups with fewer than 500 cells were excluded, and groups containing more than 10,000 cells were subsampled while preserving cluster composition.

We removed mitochondrial, ribosomal, predicted, RIKEN cDNA, sex-linked, and hemoglobin genes. Within each group, genes were further required to be detected in at least 10% of cells and in at least 50 cells. Gene expression was residualized against the number of detected genes and mitochondrial CPM fraction, then centered and scaled before calculating pairwise Pearson correlations.

To account for correlations expected by chance, we calibrated correlation thresholds separately for each group using permutation. The positive correlation cutoff was defined as the larger of 0.20 and the empirical noise threshold, and the negative cutoff as the larger of 0.30 and the empirical noise threshold. Genes without a surviving positive correlation were excluded.

Modules were identified using a signed two-layer gene graph. Positive edges connected gene pairs above the positive correlation threshold, whereas strong negative correlations were included as penalties to prevent anti-correlated genes from being assigned to the same module. We clustered this graph using the Leiden algorithm with a multiplex Reichardt-Bornholdt modularity objective and evaluated resolutions from 0.5 to 2.0, retaining the partition with the highest positive-layer modularity. Remaining within-module anti-correlated pairs were resolved by reassigning or removing conflicting genes, and modules containing fewer than 10 genes were discarded.

Large modules with weak internal correlation were recursively re-clustered at increasing Leiden resolutions. Candidate subdivisions were retained only when reproducible across two independent cell subsets, using a mean adjusted Rand index of at least 0.50. Each final module was summarized by its eigengene, defined as the first principal component of standardized residual expression. Module membership was quantified by the Pearson correlation between each gene and its module eigengene, and the 10 genes with the highest module membership were defined as hub genes.

We compared modules across cell groups based on gene-set overlap to identify recurrent transcriptional programs and quantified relationships between modules using eigengene correlations. Modules were also screened for strong associations with donor, sex, anatomical region, subclass identity, or sequencing depth. This procedure identified 3,095 modules across 265 subclasses and 1,242 modules across 125 classes by anatomical-division groups.

### Gene Ontology enrichment analysis

We performed Gene Ontology over-representation analysis using clusterProfiler^39^ v4.14.6 in R v4.4.3 with enrichGO, org.Mm.eg.db v3.20.0, and GO.db v3.20.0. The background consisted of all 25,730 GO-annotated mouse genes. P values were adjusted using the Benjamini-Hochberg procedure, and terms with adjusted P < 0.05 were considered significant.

For visualization, we focused on significant Biological Process terms supported by at least five module genes. Redundant terms were removed based on gene-set overlap using a Jaccard index threshold of 0.5, retaining the more significant term. For each module, we displayed the 12 most significant non-redundant terms. Dot position represents −log10 adjusted P value, dot size represents the number of module genes associated with the term, and dot color represents fold enrichment.

### Cell mapping between samples

Cell mapping is essential for cross-sample (one MERFISH sample to another MERFISH sample) or cross-technology (one MERFISH sample to one scRNA-seq sample or one scRNA-seq sample back to one MERFISH sample) information transfer. Given a query set *U* and a reference set *V*, our goal is to have a one-to-one mapping between the two sets so that every cell in *U* has a unique mapping target in *V* while the similarity of the mapped pairs should be as high as possible. This problem can be sequentially formed as a maximum bipartite matching problem.

For cells in the query set *U* and cells in the reference set *V*, we constructed edges *E* between all possible pair from these two sets with weight equals the proximity between the corresponding two cells measured by Predictor, resulting in a bipartite graph *G* = ((*U, V*), *E*). We wanted to find a matching *M* ⊆ *E* so that:

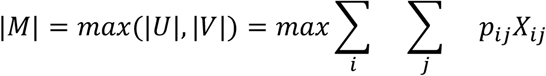

where 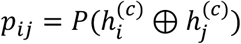 is the proximity between cell *i* in *U* and cell *j* in *V* while *X*_*ij*_ as an boolean indicator controlling if cell *j* is mapped to cell *i*. We solved the problem using the Jonker-Volgenant algorithm implemented by Scipy^40^. The matching *M* with index pairs stored in *X*_*ij*_ was then referred to transfer cell coordinates from the reference set to the query set.

For cross-sample mapping in MERFISH, we randomly selected a reference sample that is physically next to the query sample. The full reference-query table is listed in supplementary (**Supplementary Table 5**).

### Detection of cellular neighborhoods

Detection of cellular neighborhoods, local microenvironment surrounding cells, was performed on predicted spatial dissimilarity rather than the original cell-gene graph. Firstly, pairwise proximity between every possible pair of cells was predicted by Predictor. The spatial dissimilarity between cell pair *i* and *j* with embedding vector 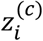 and 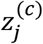 on the other hand, is defined by:

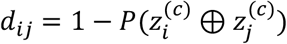

since we expected cell pairs with high proximity to have lower spatial dissimilarity and more close together. Secondly, a dissimilarity graph was constructed with an adjacency matrix *D* = (*d*_*ij*_). The graph is a dense and bidirected graph where every two nodes share an edge representing their dissimilarity. A kNN search with numbers of neighbors 15 was then conducted on top of the graph before connectivities were derived using UMAP. Thirdly, the Leiden clustering^41^ with certain resolution was applied on the connectivity graph to detect interconnected groups of nodes. Due to the differences in sample size, we set resolution as 0.5 for the WMB-10Xv3 dataset and 0.35 for AD datasets. These groups of nodes can be regarded as cellular neighborhoods which are highly connected communities spatially. All steps above were achieved using incorporated functions in Scanpy^42^.

We detected cellular neighborhoods separately in each sample (brain section from MERFISH or brain region sample from scRNA-seq) to ensure meaningful outputs. Cell type composition and ligand-receptor interaction are then calculated based on single-cell profiles in each individual cellular neighborhood.

### Ligand-receptor interaction

The ligand-receptor interaction was inferred solely from the expression matrices. For each target niche, we first selected its gene expression from the original sample by cell indices. The expression matrix was then used to infer cell-cell interaction by measuring average gene expression of proteins involved. We ran CellPhoneDB^21^ incorporated by LIANA^20^ with the mouse consensus database and 10% expression proportion cutoff and generated mean ligand-receptor expression values as well as their permutation-based p-values which are grouped by cell types with all possible (sender cell type, ligand, receptor receiver cell type) combinations. To visualize differences in ligand-receptor interaction between two niches, unique tuples of (sender cell type, ligand, receptor receiver cell type) with FDR corrected p-values < 0.05 were identified for each niche. The results were then illustrated by CCPlotR^20,43^ as interacting modes in circular plots and arrow plots.

### Differential expression within inferred cellular neighborhoods

Inferred cellular neighborhoods were identified independently within each sample by Leiden clustering of the ProxiNet-derived cell proximity graph using the same parameters for WT and 5xFAD samples (resolution 0.35, k=15 nearest neighbors, random seed 0). Differential expression analysis focused on the 8-month 5xFAD entorhinal cortex sample, which contained 5,011 nuclei. Only nuclei with confidently assigned major cell-type labels were included.

For each major cell type, we compared nuclei assigned to neighborhood AD0 with nuclei of the same cell type in all other 8-month 5xFAD neighborhoods. We required at least 20 nuclei in both groups, allowing analysis of GABAergic neurons (229 AD0 vs 552 other), astrocytes (215 vs 214), oligodendrocytes (184 vs 60), and microglia (122 vs 93). OPCs, glutamatergic neurons, and endothelial cells did not meet this criterion and were excluded from differential expression testing. Genes were retained if they were detected in at least 10% of nuclei in either comparison group.

We assessed differential expression using two complementary tests and retained only genes supported by both. First, we fitted a negative-binomial generalized linear model to raw UMI counts, including library size as an offset and sequencing depth, mitochondrial fraction, and ribosomal fraction as covariates. Second, we applied a two-sided Wilcoxon rank-sum test to library-size-normalized expression. P values from each test were adjusted separately using the Benjamini-Hochberg procedure. Genes were considered differentially expressed when they reached FDR < 0.05 in both tests, showed concordant direction of change, and had an absolute log2 fold change of at least 0.25.

We performed several control analyses to assess robustness. We repeated the analysis after permuting AD0 labels within each cell type and observed no consensus differential expression calls under permutation. We also repeated the comparisons after matching cells between groups by sequencing depth to evaluate sensitivity to residual library-size differences. Because long, intron-rich genes can be overrepresented in single-nucleus RNA-seq, we compared gene-length distributions between differentially expressed genes and the tested background. Potential signals driven by cell-type contamination were further examined using marker expression and doublet scores. Because only one library was available per genotype, we did not perform statistical inference between WT and 5xFAD samples; WT comparisons were used descriptively only.

## Data availability

All datasets analyzed in this study were previously published and are publicly available. The whole-mouse-brain MERFISH dataset was obtained from Zhang et al.^11^ (https://cellxgene.cziscience.com/collections/0cca8620-8dee-45d0-aef5-23f032a5cf09). The STARmap dataset was obtained from Shi et al.^12^ (https://doi.org/10.5281/zenodo.8327576). The Slide-seq dataset was obtained from Langlieb et al.^13^ (www.BrainCellData.org). The WMB-10Xv3 single-cell RNA-seq dataset was obtained from Yao et al.^22^ (https://github.com/AllenInstitute/abc_atlas_access/blob/main/descriptions/WMB-10Xv3.md). The 2-month and 8-month entorhinal cortex snRNA-seq datasets used for the Alzheimer’s disease analysis were obtained from Bartas et al.^23^ (GSE296091).

The train, validation, and test sample assignments used in this study are provided in Supplementary Tables 1-3. Additional processed data supporting the analyses will be deposited at Zenodo upon publication. No new raw sequencing or spatial transcriptomic data were generated in this study.

## Code availability

The ProxiNet source code, together with the scripts and Jupyter notebooks used for model training, evaluation, and generation of the results presented in this study, is available at the ProxiNet GitHub repository https://github.com/nasunlab/ProxiNet. The version of the code corresponding to this manuscript will be archived in Zenodo with a persistent DOI. Pretrained ProxiNet models, including model parameters for cellular-proximity prediction and auxiliary coordinate-inference analyses, will also be deposited in the same Zenodo repository.

## Acknowledgements

This work was supported by the Whitehead Institute and the Whitehead Innovation Initiative. N.S. was supported by the Whitehead Institute Fellows Program and the NIH Director’s Early Independence Award (DP5OD042638). The content is solely the responsibility of the authors and does not necessarily represent the official views of the NIH or other funding agencies.

## Author contributions

N.S. conceived and designed the study and supervised the project. B.Y. developed the initial version of the model. Y.Z. developed the computational framework and performed the data analyses with assistance from A.Z.. M.K. provided scientific input. Y.Z., A.Z., and N.S. wrote and revised the manuscript with input from all authors.

## Supplementary Figures

**Supplementary Figure 1.**
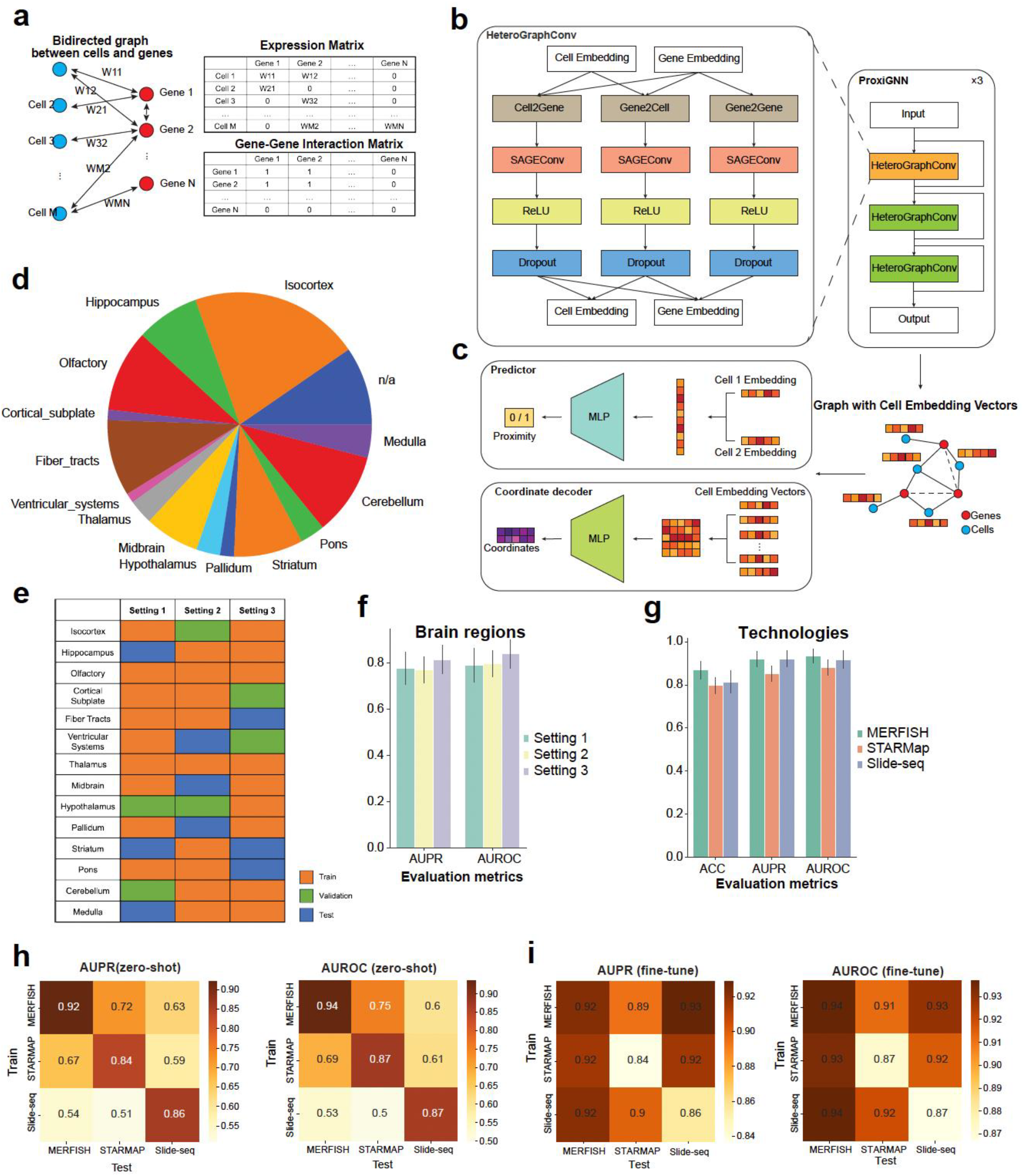
Detailed ProxiNet architecture and evaluation across brain regions and spatial technologies. **a**, Construction of the heterogeneous cell–gene graph used by ProxiNet. The expression matrix defines weighted bidirectional edges between cell and gene nodes, and the gene–gene interaction matrix defines prior-informed edges among gene nodes. **b**, Detailed architecture of ProxiGNN. Relation-specific message passing is performed separately for Cell2Gene, Gene2Cell, and Gene2Gene relations using SAGEConv layers followed by ReLU activation and dropout. Three sequential HeteroGraphConv layers generate updated cell and gene embeddings. **c**, Downstream use of learned cell embeddings. For the primary proximity-prediction task, embeddings from two cells are combined and passed to an MLP predictor to estimate binary cellular proximity. An auxiliary coordinate decoder maps learned cell embeddings to spatial coordinates to assess whether spatial information is retained in the learned representation. **d**, Representation of major mouse brain regions included in the spatial transcriptomic datasets used for model development and evaluation. **e**, Three brain-region train, validation, and test configurations used to evaluate generalization to held-out anatomical regions. **f**, Proximity-prediction performance across the three brain-region evaluation settings, quantified by AUPR and AUROC. **g**, Proximity-prediction performance across MERFISH, STARmap, and Slide-seq datasets, quantified by accuracy, AUPR, and AUROC. Bars show evaluation metrics and error bars indicate standard deviation across test sections. **h**, Zero-shot prediction across technology. Heat maps show AUPR and AUROC when models trained on MERFISH, STARmap, or Slide-seq are evaluated on each of the three spatial technologies. **i**, Fine-tune prediction across technology. Heat maps show AUPR and AUROC when models trained on MERFISH, STARmap, or Slide-seq are finetuned and evaluated on each of the three spatial technologies.

**Supplementary Figure 2.**
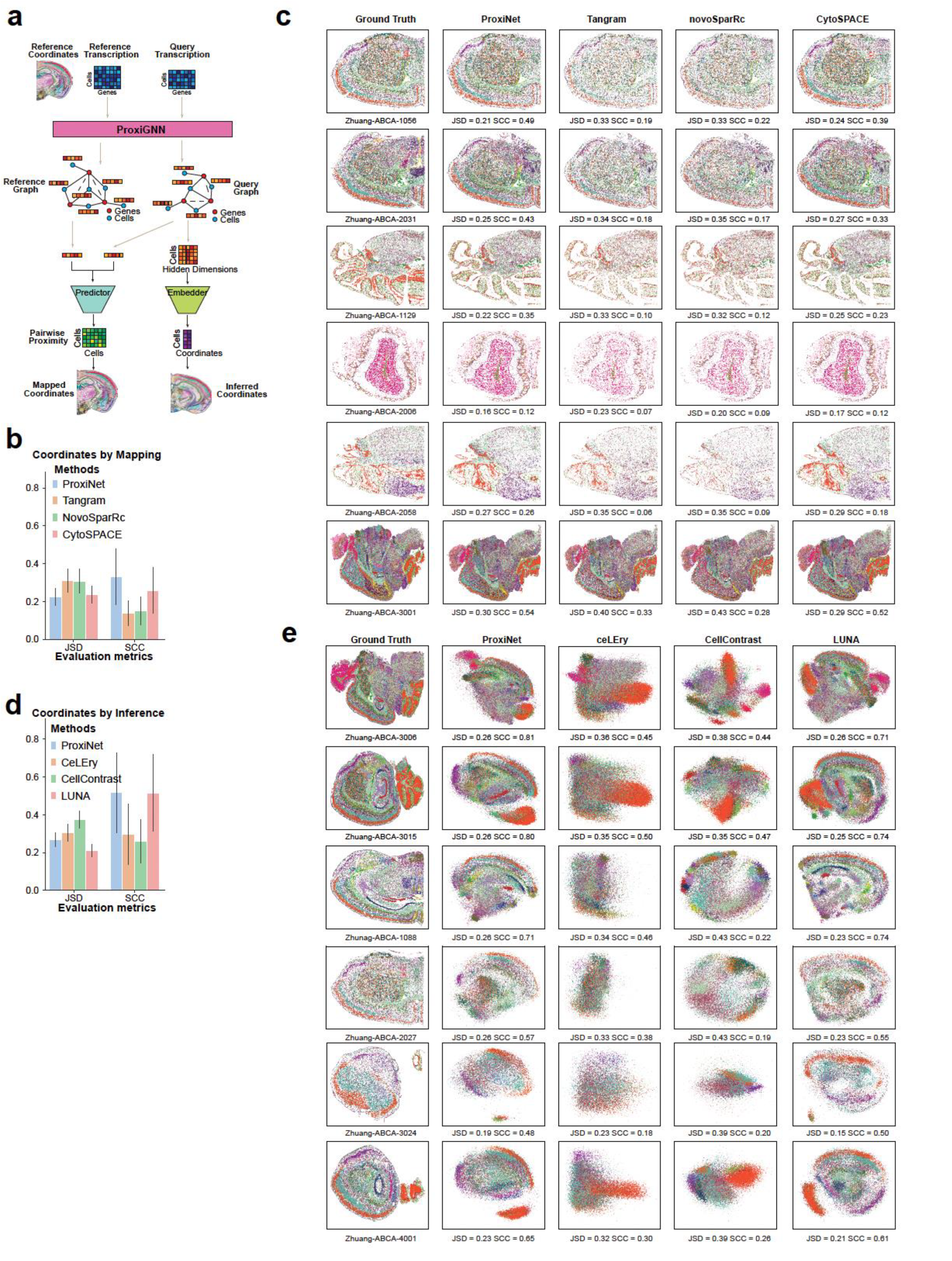
ProxiNet representations retain spatial information in auxiliary coordinate-mapping and inference tasks. **a**, Overview of coordinate mapping and coordinate inference using ProxiNet representations. For coordinate mapping, reference spatial coordinates and transcriptomes are used together with query transcriptomes to estimate pairwise relationships and map query cells onto the reference spatial organization. For coordinate inference, learned cell representations are passed through a coordinate decoder to predict spatial coordinates directly from transcriptomic information. **b**, Quantitative comparison of coordinate mapping using ProxiNet, Tangram, NovoSpaRc, and CytoSPACE, evaluated by Jensen–Shannon divergence (JSD) and Spearman correlation coefficient (SCC). Lower JSD and higher SCC indicate greater agreement with ground-truth spatial organization. **c**, Representative coordinate-mapping results across Zhuang-ABCA-1056, Zhuang-ABCA-2031, Zhuang-ABCA-1129, Zhuang-ABCA-2006, Zhuang-ABCA-2058, and Zhuang-ABCA-3001. Ground-truth spatial organization is shown alongside results from ProxiNet and comparator methods; JSD and SCC values are indicated for each reconstruction. **d**, Quantitative comparison of coordinate inference using ProxiNet, CeLEry, CellContrast, and LUNA, evaluated by JSD and SCC. **e**, Representative coordinate-inference results across Zhuang-ABCA-3006, Zhuang-ABCA-3015, Zhuang-ABCA-1088, Zhuang-ABCA-2027, Zhuang-ABCA-3024, and Zhuang-ABCA-4001. Ground-truth spatial organization is shown alongside inferred coordinates from each method, with JSD and SCC values indicated.

**Supplementary Figure 3.**
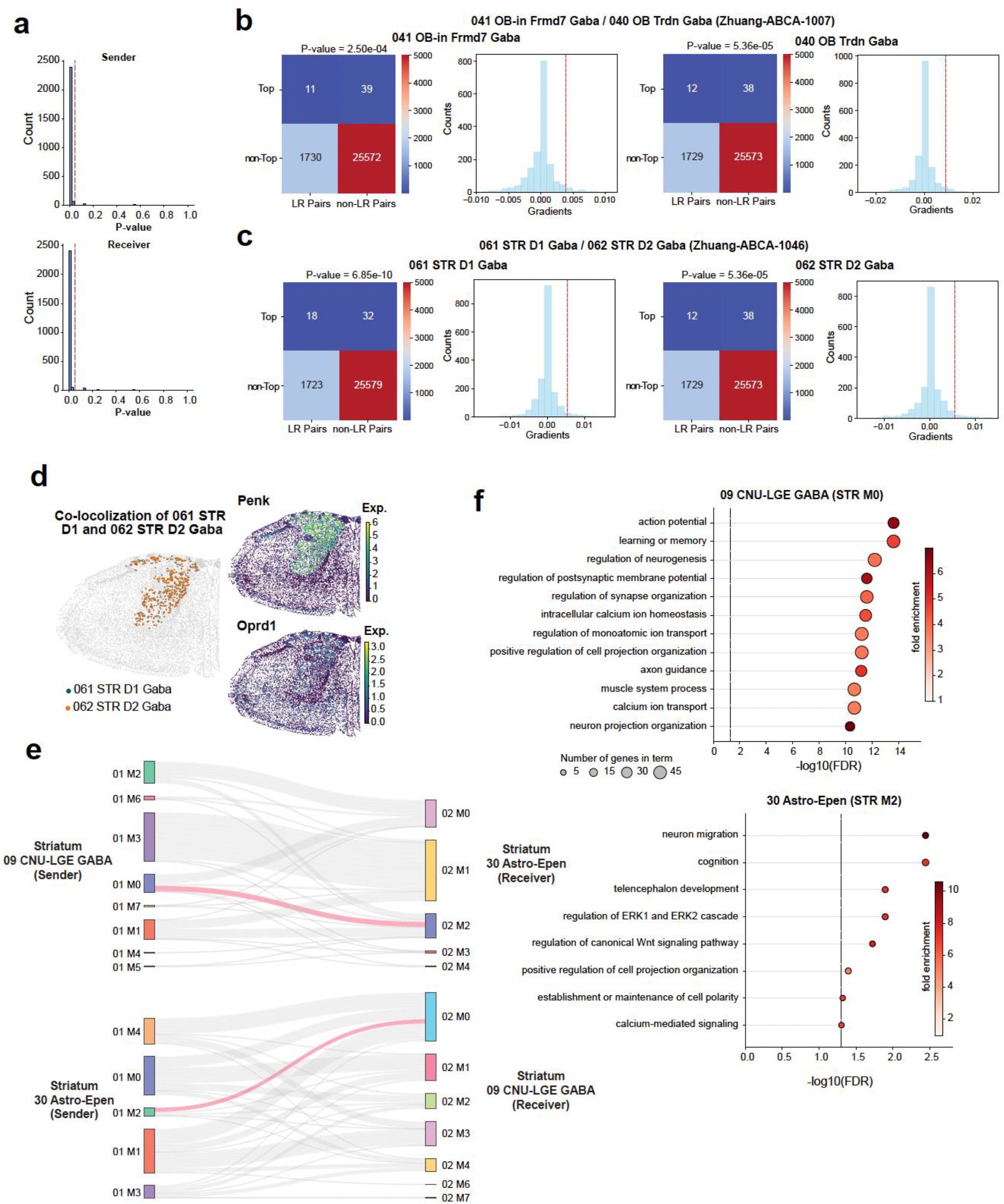
Proximity-associated genes are enriched for ligand–receptor relationships and nominate broader transcriptomic programs. **a**, Distribution of significance values for sender- and receiver-associated genes identified by gradient-based attribution across evaluated cell-type pairs. **b**, Enrichment of ligand–receptor genes among highly attributed genes for the 041 OB-in Frmd7 GABA and 040 OB Trdn GABA cell populations in Zhuang-ABCA-1007. Contingency tables compare ligand–receptor (LR) and non-LR genes among top versus non-top attribution-ranked genes, with enrichment P values shown above each comparison. Histograms show the corresponding distributions of gradient-based attribution scores, with dashed lines indicating the selected attribution cutoff. **c**, Equivalent ligand–receptor enrichment analysis for 061 STR D1 GABA and 062 STR D2 GABA cells in Zhuang-ABCA-1046. **d**, Spatial co-localization of STR D1 GABA and STR D2 GABA populations and spatial expression of Penk and Oprd1, providing an independent example of a ligand– receptor pair prioritized through ProxiNet attribution. **e**, Relationships between proximity-associated transcriptomic modules in Striatum 09 CNU-LGE GABA and 30 Astro-Epen cells in both sender–receiver directions. Highlighted connections denote ligand–receptor relationships prioritized between the indicated modules. **f**, Gene Ontology enrichment of representative proximity-associated modules in Striatum 09 CNU-LGE GABA (M0) and 30 Astro-Epen (M2) cells. Dot position denotes −log10FDR, dot color denotes fold enrichment, and dot size denotes the number of genes assigned to each term.

**Supplementary Figure 4.**
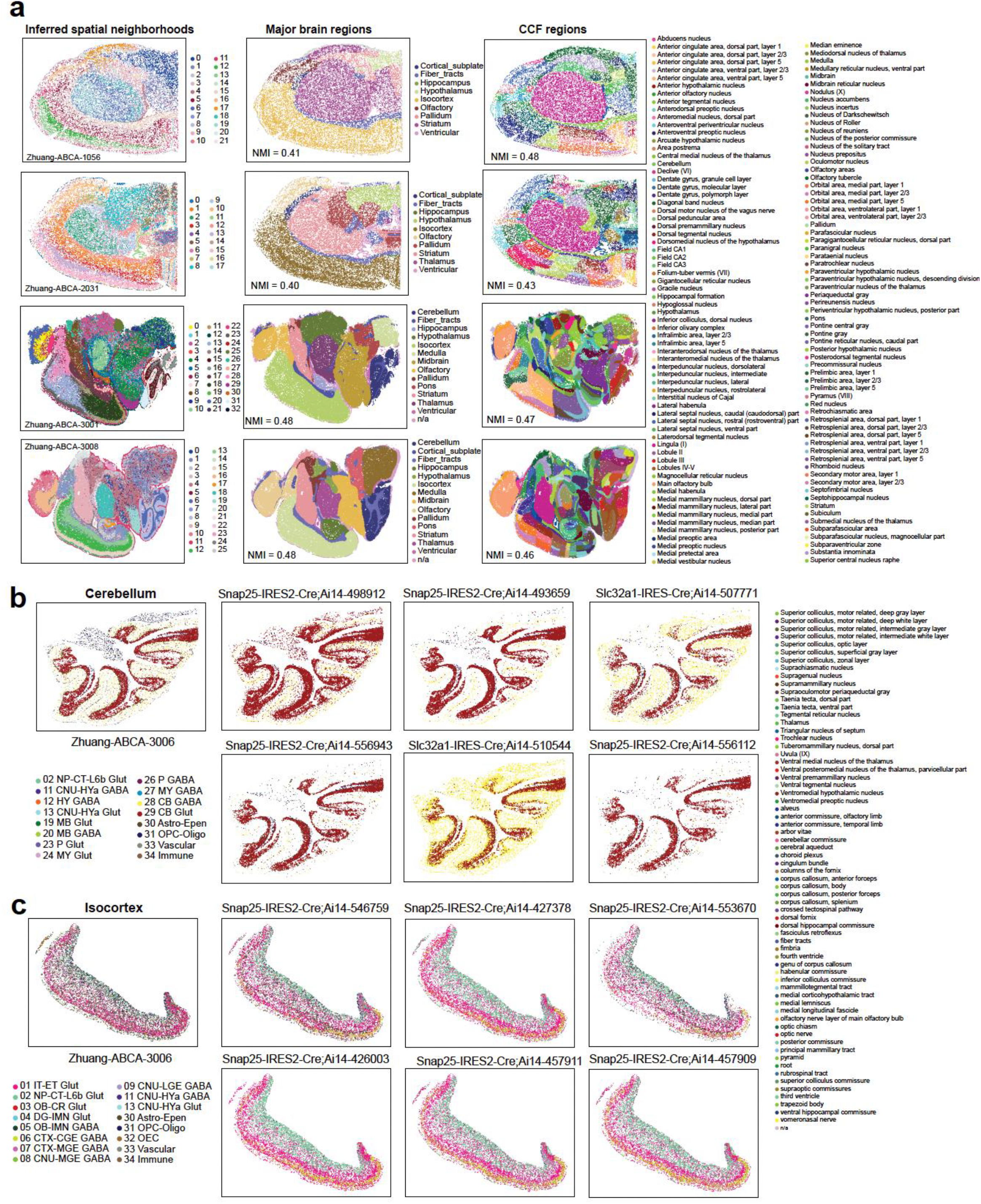
Inferred cellular neighborhoods reproducibly recover anatomical tissue organization across spatial datasets. **a**, Expanded comparison of inferred cellular neighborhoods with anatomical organization across four spatial transcriptomic sections: Zhuang-ABCA-1056, Zhuang-ABCA-2031, Zhuang-ABCA-3001, and Zhuang-ABCA-3008. For each section, cells are colored by inferred neighborhood assignment, annotated major brain region, or Common Coordinate Framework (CCF) region. NMI values quantify concordance between inferred neighborhoods and the corresponding anatomical annotations. **b**, Cell-type organization in the cerebellum across independent spatial transcriptomic samples. Zhuang-ABCA-3006 is shown as a reference together with multiple independent sections from the indicated mouse lines; cells are colored by major cell type to illustrate reproducibility of cerebellar cellular organization across samples. **c**, Equivalent comparison across independent Isocortex spatial transcriptomic samples, showing reproducible organization of major neuronal and non-neuronal cell populations across sections.

**Supplementary Figure 5.**
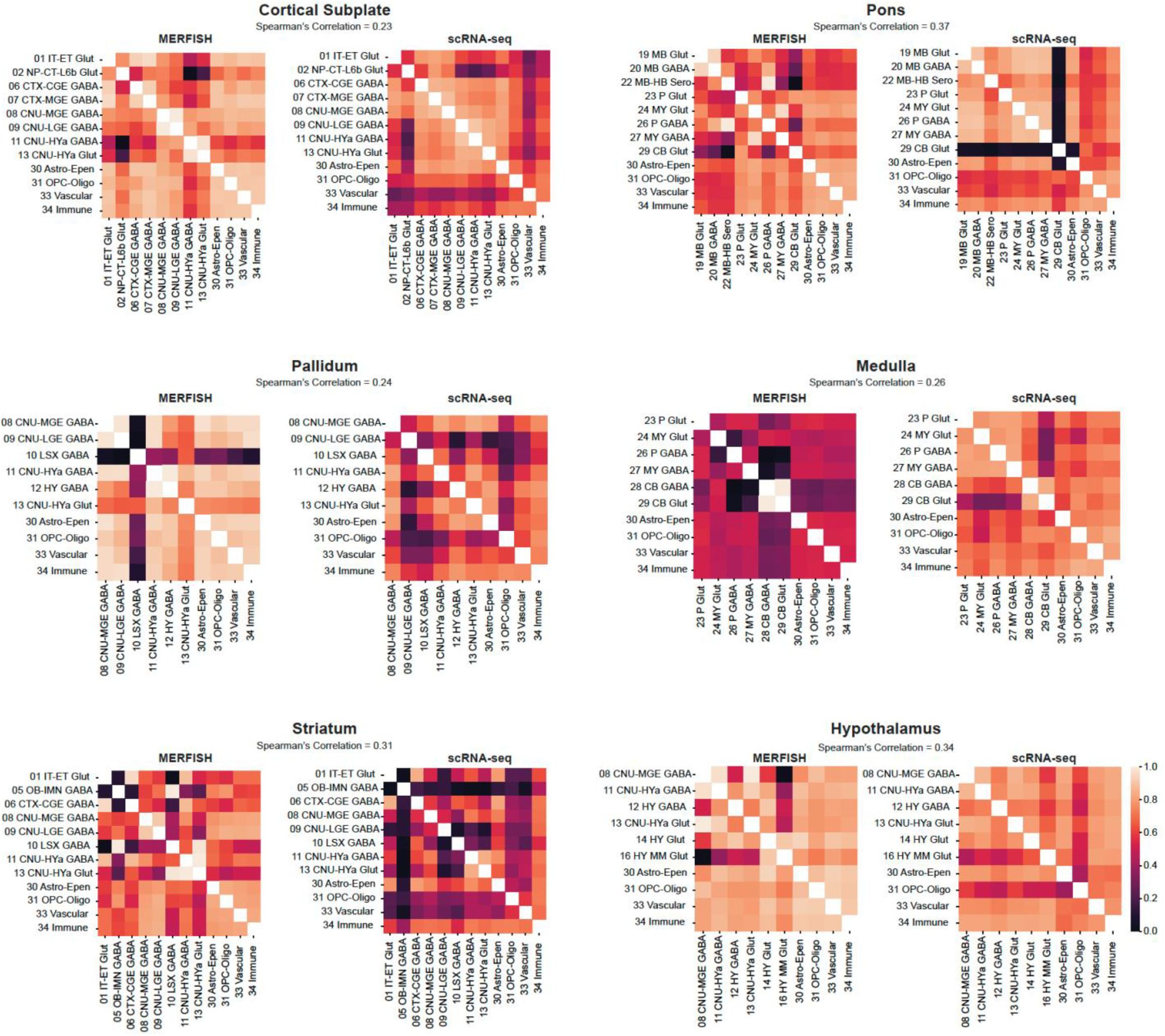
ProxiNet recovers cell-type proximity relationships across multiple brain regions in dissociated scRNA-seq. Comparison of cell-type proximity relationships measured directly in MERFISH with those inferred by ProxiNet from dissociated scRNA-seq across additional brain regions. For each region, the left heat map shows MERFISH-derived cell-type proximity relationships, and the right heat map shows the corresponding relationships inferred from scRNA-seq using the spatially calibrated ProxiNet model. Concordance between the two matrices was quantified using the Spearman correlation coefficient (SCC): Cortical Subplate, SCC = 0.23; Pons, SCC = 0.37; Pallidum, SCC = 0.24; Medulla, SCC = 0.26; Striatum, SCC = 0.31; and Hypothalamus, SCC = 0.34. Together with the Cerebellum and Olfactory examples shown in Fig. 4, these analyses evaluate transfer of spatially learned cellular-proximity relationships across anatomically distinct brain regions.

**Supplementary Figure 6.**
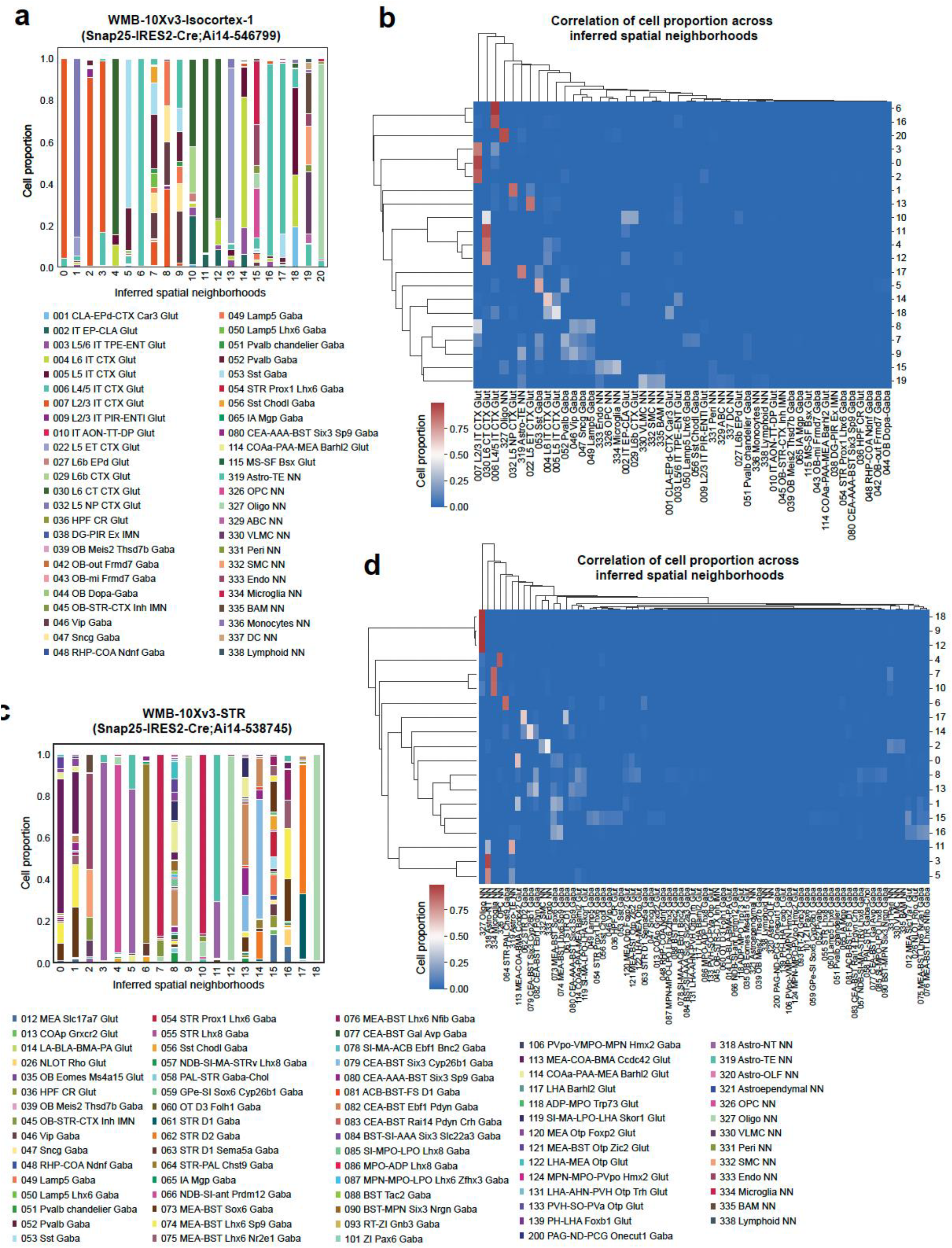
Inferred cellular neighborhoods capture fine-grained cell-subclass composition in Isocortex and Striatum. **a**, Fine-grained cell-subclass composition of inferred cellular neighborhoods in the WMB-10Xv3-Isocortex-1 scRNA-seq sample (Snap25-IRES2-Cre;Ai14-546799). Each stacked bar represents the subclass composition of one inferred neighborhood. **b**, Hierarchically clustered representation of cell-subclass proportions across Isocortex neighborhoods. Rows correspond to inferred neighborhoods, columns correspond to annotated cell subclasses, and color denotes the proportion of each subclass within a neighborhood. Dendrograms group neighborhoods and subclasses according to similarity in composition. **c**, Fine-grained cell-subclass composition of inferred cellular neighborhoods in the WMB-10Xv3-STR scRNA-seq sample (Snap25-IRES2-Cre;Ai14-538745). **d**, Hierarchically clustered representation of cell-subclass proportions across Striatum neighborhoods, showing heterogeneous combinations of neuronal and non-neuronal subclasses across inferred cellular neighborhoods.

**Supplementary Figure 7.**
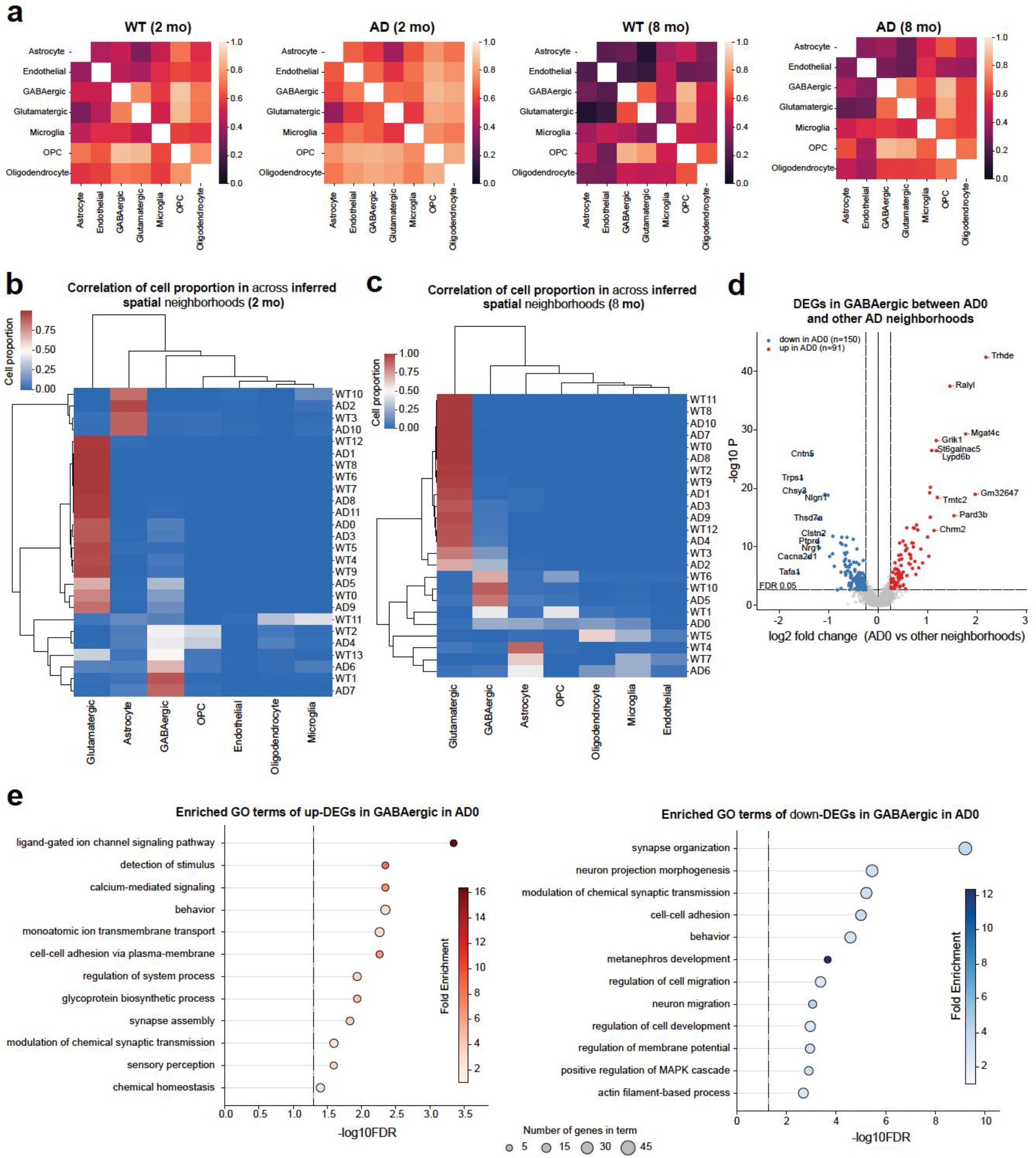
Expanded characterization of age-dependent Alzheimer’s-associated remodeling of inferred cellular neighborhoods. **a**, Predicted pairwise cell-type proximity relationships in entorhinal cortex from 2-month WT, 2-month AD, 8-month WT, and 8-month AD samples. Heat maps summarize ProxiNet-inferred proximity relationships among glutamatergic neurons, GABAergic neurons, astrocytes, oligodendrocytes, OPCs, microglia, and endothelial cells. **b**, Hierarchically clustered cell-type composition of inferred cellular neighborhoods at 2 months across WT and AD samples. Rows represent individual neighborhoods and columns represent major cell types; color denotes cell-type proportion. **c**, Equivalent analysis of inferred cellular neighborhoods at 8 months, illustrating changes in neighborhood composition and WT–AD correspondence with disease progression. **d**, Differential gene-expression analysis of GABAergic neurons in AD0 versus GABAergic neurons from other AD neighborhoods at 8 months. The volcano plot shows log2 fold change and −log10P; 91 genes were upregulated and 150 were downregulated in AD0 at FDR < 0.05. Selected genes are labeled. **e**, Gene Ontology enrichment of genes upregulated (left) and downregulated (right) in AD0 GABAergic neurons. Dot position denotes −log10FDR, dot color denotes fold enrichment, and dot size denotes the number of genes assigned to each term.

